# Complete timing of copy-number amplification histories reveals whole-genome duplication dynamics and coincident oncogene amplification bursts

**DOI:** 10.64898/2026.09.23.753832

**Authors:** Joseph Brew, Peter J. Park, Doga C. Gulhan

**Affiliations:** Department of Biomedical Informatics, Harvard Medical School, Boston, MA, USA; Krantz Family Center for Cancer Research, Mass General Brigham, Boston, MA, USA; Department of Medicine, Harvard Medical School, Boston, MA, USA; Broad Institute of MIT and Harvard, Cambridge, MA, USA

## Abstract

Copy-number amplifications drive cancer evolution through increased oncogene dosage. Mutations acquired before a gain are co-amplified, so their multiplicity places successive gains on a molecular timescale. This principle has been used to time individual gains and whole-genome duplication (WGD), but possible gain histories multiply rapidly with copy number, confining existing methods to low copy-number states or predefined scenarios and leaving oncogene-bearing amplifications unresolved. Here we describe a mathematical framework to systematically enumerate every allele-specific gain route to a copy-number state and derive an analytical timing for each gain, implemented in a method called Tau. Across pan-cancer genomes, punctuated multichromosomal events account for roughly three-quarters of clonal gains. Most are WGD, acquired early and often repeated within a doubling-prone subset, but a distinct, predominantly monoallelic class recurs on tumor-type-specific chromosomes, representing a coordinated mode separate from doubling. Strikingly, the highest copy-number states are reached not gradually but in abrupt bursts that cluster around WGD and contain more gains than doubling alone can produce, recurring at oncogenes. WGD is therefore not a passive doubling but a window in which localized oncogene amplification becomes far more frequent. By resolving the order of gains and the mutational processes shifting between them, Tau establishes punctuated amplification as a pervasive mode of cancer genome evolution.

## Introduction

During cancer evolution, genomic alterations that confer selective advantages accumulate within expanding cellular populations. Copy-number alterations are among the most prevalent changes in cancer genomes and contribute to tumor initiation, progression, treatment resistance, and intratumor heterogeneity; extensive chromosomal instability is also associated with adverse clinical outcomes [1–3].

Genome-wide and multichromosomal gains can arise through whole-genome duplication (WGD), chromosome mis-segregation, or complex rearrangement processes affecting multiple chromosomes [4–6]. Focal amplification mechanisms can further increase copy number at individual genomic loci [7–9]. Together, these processes can generate high-copy-number states, particularly at oncogenic loci, increasing gene dosage and, in some contexts, reorganizing chromatin and regulatory interactions to enhance oncogene transcription [10]. Distinct mechanisms can nevertheless converge on similar observed copy-number states while producing markedly different temporal patterns of gain, from gradual accumulation to punctuated amplification affecting individual loci, multiple chromosomes, or the entire genome. Resolving the timing of successive copy-number gains is therefore essential for reconstructing how amplifications arise and shape subsequent tumor evolution.

Advances in cancer whole-genome sequencing have enabled increasingly detailed reconstruction of tumor evolutionary histories [11, 12]. Clonal reconstruction and multi-sample phylogenetic approaches can identify tumor sub-clones and infer their evolutionary relationships [13–15]. However, copy-number-based reconstruction from bulk sequencing remains challenging because subclonal copy-number alterations are difficult to detect and quantify [13, 15]. Single-cell sequencing can resolve these alterations more directly [16, 17], but typically involves a trade-off between the number of cells profiled and the depth of mutational characterization within each cell [18, 19]. Moreover, although phylogenetic approaches can assign events to subclonal branches that diverged after the most recent common ancestor (MRCA), events acquired along the shared clonal lineage leading to the MRCA generally remain collapsed onto a single unresolved trunk. Copy-number timing methods recover information from this earlier history – up until the MRCA time point – by exploiting a simple observation: mutations present on an allele before it is gained are co-amplified and therefore reach higher multiplicity than mutations acquired afterward [11, 20, 21]. The multiplicity distribution of somatic mutations can thus place gains on a mutation-based molecular timescale. Because mutation rates can change during tumor evolution, molecular time does not map linearly onto chronological time [11]; nevertheless, it can reveal ordering and synchrony of events across the genome.

The same observed high-copy-number state can arise through many distinct gain histories, making those histories increasingly difficult to reconstruct as copy number rises. Existing methods have established the value of molecular timing but differ in scope. MutationTimeR’s framework resolves gains with WGD-associated histories but up to limited copy-number values [11]; Butte estimates the intervals before the first and after the last gain up to high copy-number states [22]; AmplificationTimeR infers parsimonious orders for regions shaped by sequential gains [23]; GRITIC extended timing to higher-copy number gains and non-parsimonious solutions in tumors with a single genome duplication or no duplication [24]. As allele-specific copy number increases, however, the number of distinct gain histories (or routes) compatible with the same observed major and minor copy-number state grows rapidly. Existing models manage this complexity with simplifying assumptions, solve only a restricted set of copy-number states, or impose conditions on synchronicity of the gains through WGD to simplify the problem. No existing approach systematically enumerates all allele-specific gain routes across arbitrary copy-number states and analytically characterizes exact or bounded timings for every successive gain.

To address these limitations, we developed Tau (Timing Amplifications under Uncertainty), a generalized analytical framework that extends copy-number timing beyond predefined evolutionary scenarios. Tau systematically enumerates evolutionary routes from a diploid ancestor to arbitrary allele-specific copy-number states and derives exact timing solutions or constrained solution spaces for successive gains (**Figure 1a–c**). Precomputing these solutions makes exhaustive evaluation of the evolutionary gain route space computationally tractable. By integrating amplification histories across genomic segments, Tau groups compatible routes and identifies synchronous gains affecting different chromosomes and alleles, enabling the detection of WGD and other cross-chromosomal punctuated events without imposing their presence beforehand. Tau provides a framework for reconstructing complex amplification histories and for investigating the temporal organization of copy-number evolution across cancer genomes.

**Figure 1:**
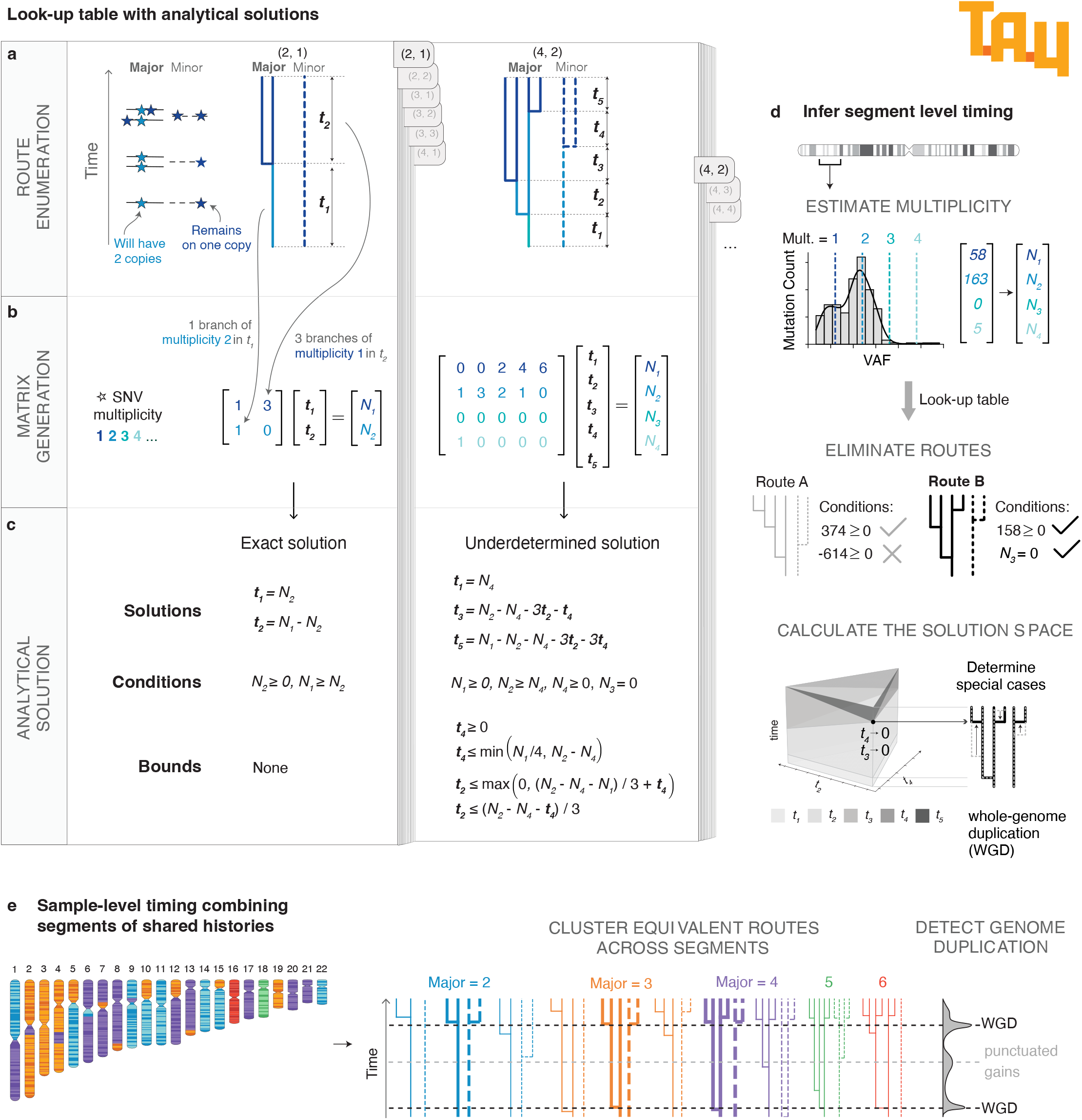
Tau enumerates all gain routes to a copy-number state and solves their timing analytically. **a**, Construction of the look-up table, illustrated for a determined state (2, 1) and an underdetermined state (4, 2). Top, all allele-specific routes from a diploid ancestor are enumerated, each a sequence of gains separated by intervals of molecular time. **b**, Mutations present before a gain are propagated to the resulting copies, so each route defines a linear system relating the intervals {*t*_*i*_} to the SNV multiplicity counts {*N*_*i*_}, with one column per interval and one row per multiplicity. **c**, Each system is solved analytically, yielding equalities for {*t*_*i*_}, conditions on {*N*_*i*_} that define the route’s feasible region, and bounds on {*t*_*i*_}. Determined routes give a unique solution; underdetermined routes give a bounded solution space. **d**, Segment-level inference. Multiplicity counts are estimated from the variant allele fraction distribution, routes whose conditions the counts violate are eliminated, and the retained route’s solution space is evaluated. Special cases in which gains coincide, such as whole-genome duplication (WGD), are identified within that space rather than assumed. **e**, Sample-level integration. Segments sharing a copy-number state and route are pooled and re-timed, and their gain times are clustered across the genome to detect punctuated gains shared by multiple chromosomes, including WGD.

Applied to the Pan-Cancer Analysis of Whole Genomes (PCAWG) cohort, Tau distinguished two common classes of large-scale, multichromosomal gain events: WGD and predominantly monoallelic gains affecting tumor-type-specific sets of chromosomes. By resolving successive WGDs, Tau revealed their temporal dynamics and substantial interpatient heterogeneity in the propensity to undergo genome doubling. In high-copy-number regions, Tau further uncovered localized oncogene-amplification bursts that clustered around WGD and comprised more gains than doubling alone could explain. Together, these analyses provide a high-resolution temporal reconstruction of copy-number evolution that distinguishes coordinated gain modes, resolves the dynamics of repeated WGD, and reveals a previously unresolved temporal association between WGD and localized oncogene amplification.

## Results

### Tau provides a generalized framework for timing any copy-number amplification

Tau was designed to address a central challenge in copy-number timing: as the major and minor copy numbers increase, the number of amplification routes compatible with the observed state grows rapidly. For instance, a state of (Major CN, Minor CN) = (3, 2) has two mathematically distinct routes, while a state of (7, 5) has 6009 (**Supplementary Figure S1a**, Methods). Existing approaches therefore tend to focus on specific classes of amplifications or predefined evolutionary scenarios such as WGD that constrain both alleles to be gained at the same time [11, 23, 24]. Tau instead systematically enumerates all valid gain routes from an ancestral diploid state to a target copy-number state (**Figure 1a**). Each route represents a sequence of amplification events separated by intervals of molecular time (*t*_1_, *t*_2_, …), and the propagation of mutations on amplified segments defines a linear relationship between these intervals and the observed mutation multiplicity distribution. As gains accrue, SNVs acquired in different time intervals, or on different branches, are propagated to different numbers of copies and so appear at different multiplicities. This relationship can be represented as a matrix, each column corresponding to a time interval (*t*_1_, *t*_2_, …) and each row a SNV multiplicity (*N*_1_, *N*_2_, …) (**Figure 1b**). The sets of linear equations defined by these matrices are solved analytically with SageMath [25].

Depending on its identifiability, a route yields either an exact timing solution (determined) or a constrained solution space (underdetermined) (**Figure 1c**), allowing incompatible histories to be excluded and the timing of gains along compatible routes to be estimated or bounded (see Methods). In the case of underdetermined solutions, the constrained solution space obtained corresponds to an infinite set of equally admissible solutions satisfying the constraints, and that cannot be distinguished from each other without further assumptions. To make the evaluation of every route computationally tractable, we generated routes recursively (**Supplementary Figure S1b**), solved each route’s system of equations analytically in advance, and stored the resulting timing solutions in a look-up table. Pre-computed solutions currently cover copy-number states up to (7, 5) and can be extended further if needed. Together, these states in the look-up table of analytical solutions cover 99.7% of the amplified genome in the PCAWG cohort. The look-up table enables nearly instantaneous solution of timing per segment.

To apply Tau to a cancer genome, we begin by timing each amplified segment (**Figure 1d**). Tau estimates the counts of mutation multiplicities (*N*_1_ =the number of mutations of multiplicity 1, and so on) from the distribution of variant allele fractions (VAFs) and substitutes these values into the corresponding precomputed analytical solutions—obtaining exact timings for determined routes or bounded intervals for underdetermined ones—and applies the accompanying constraints to exclude routes incompatible with the observed data. By eliminating incompatible routes and the need for a computationally intensive probabilistic sampling strategy used by other algorithms [24], the analytical framework of Tau substantially reduces runtime. Tau then integrates segment-level results across the genome to detect punctuated gains shared by multiple segments across chromosomes (**Figure 1e**). This integration connects individual segment histories to coherent, sample-level trajectories of genome evolution. Mutational signatures are characteristic patterns of somatic mutations that provide readouts of underlying mutational processes [26–28]. For inferring the molecular time when gains take place, Tau uses mutations attributed to the ubiquitous clock-like signatures SBS1 and SBS5 rather than the full mutation set [29], because the activities of other mutational processes can vary even more substantially during tumor evolution owing to changes in exogenous exposure or the onset of genomic instability.

Together, these components enable high-resolution, sample-level reconstruction of complex amplification histories without prespecifying a gain scenario, while explicitly retaining uncertainty when the data do not identify a unique route.

### Integrating segment histories yields high-confidence sample-level amplification trajectories

To reconstruct sample-level trajectories, Tau first times individual segments and then groups segments with the same copy-number state and compatible gain routes, considering route topology and the timing of individual gains as reflected in their multiplicity profiles (**Supplementary Figure S2**). Tau pools these grouped segments’ mutation multiplicity distributions and re-estimates the timing of successive gains, improving precision for gain events extending across multiple copy-number segments. Pooling substantially increased the number of informative mutations available for timing: 76.4% of PCAWG segments belonged to route clusters containing more than 200 SBS1/SBS5-weighted mutations, whereas only 3.7% of individual segments exceeded this threshold before pooling (**Supplementary Figure S3c**). In simulations, timing estimates based on more than 200 mutations had a median error of 0.013 molecular-time units (**Supplementary Figure S3d**).

To illustrate the resulting segment- and sample-level trajectories, we selected a stomach adenocarcinoma and an esophageal adenocarcinoma exhibiting distinct forms of punctuated amplifications (**Figure 2**). A browsable catalog of Tau timing reconstructions for all PCAWG samples, including these examples, is available at https://tau.sigscape.org/. In the stomach tumor (**Figure 2a**), gains at two time points affected both alleles across more than two-thirds of chromosomes, consistent with sequential WGDs. In the esophageal tumor (**Figure 2b**), alongside a late WGD, Tau resolved an earlier, predominantly monoallelic event affecting chromosomes 3 and 15, with smaller contributions from other chromosomes. Across PCAWG, we identified analogous punctuated events involving coordinated gains across multiple chromosomes that did not meet the criteria for WGD classification (Methods); we term these cross-chromosomal punctuated gains (ccPGs). Together with additional chromosome-specific gains, these events generated major-allele copy-number states as high as 5–7 in these examples.

**Figure 2:**
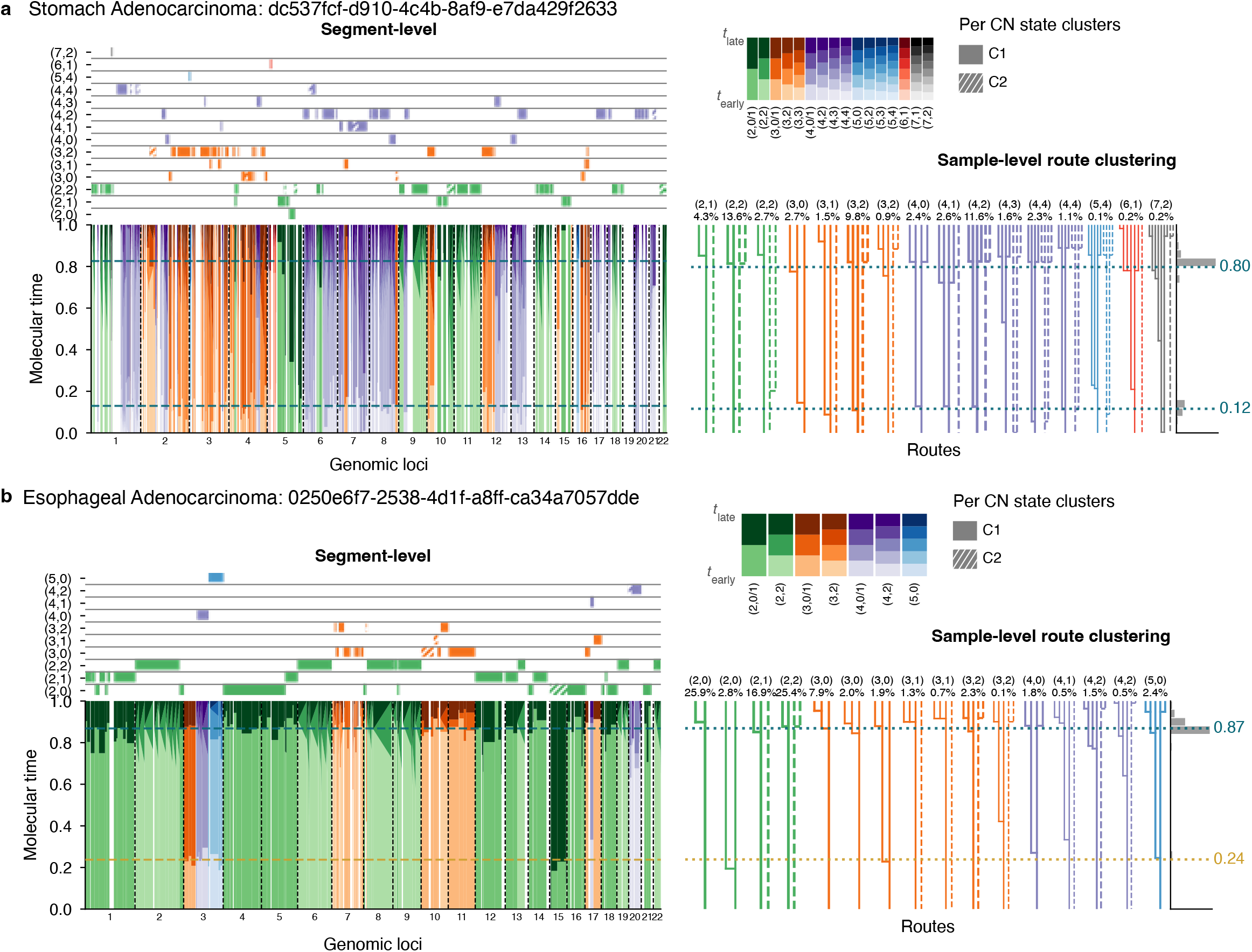
Segment timing and route clustering reveal diverse amplification trajectories converging at shared gain times. **a**, Stomach adenocarcinoma with two sequential whole-genome duplications (WGDs). Left, segment-level timing: tracks indicate timed segments of each allele-specific copy-number state along the genome, and the stacked plot below shows the inferred gain times of every segment against molecular time (*t*_early_ = 0 to *t*_late_ = 1), colored by copy-number state and within-state cluster. The marginal histogram gives the distribution of gain times across the genome; dashed lines mark the sample-level events detected by time-point clustering. Right, pooled-segment route clustering: segments sharing a copy-number state and multiplicity distribution are pooled and re-timed, and the resulting allele trees are shown for each cluster, labeled by copy-number state and percentage of the timed genome (with more than 10 SBS1/SBS5-weighted mutations including both diploid and aneuploid segments). Solid and dashed branches denote the major and minor allele. **b**, As in **a**, for an esophageal adenocarcinoma with one WGD (*t* = 0.87) and one earlier cross-chromosomal punctuated gain (*t* = 0.24) predominantly affecting chromosomes 3 and 15.

The two illustrative tumors also demonstrate why inferring unrestricted gain routes before identifying synchronized events is important. Tau derives the unrestricted solution space before identifying synchronized configurations that satisfy the analytical constraints. It can therefore recover coordinated biallelic gains, or gains affecting multiple copies of the same allele, without imposing WGD or another predefined evolutionary model a priori. For instance, in the stomach cancer example, two (2,2) pooled-segment clusters show distinct histories: one (13.6% of the genome) is consistent with a biallelic gain at the later WGD time point (**Figure 2a**, right panel second route from left), whereas the other (2.7% of the genome) is consistent with monoallelic gains at both the earlier and later WGD time points (third route from left) – a pattern that would not have been distinguished had simultaneous gains on both alleles coinciding with a WGD event been assumed as in previous methods [11, 23].

Tau reconstructs evolutionary histories at high temporal resolution while preserving uncertainty when the data do not identify a unique history. These examples show how pooling compatible segment histories improves timing precision while retaining distinct allele-specific routes within a coherent sample-level evolutionary trajectory.

### Tau accurately estimates gain times across both determined and underdetermined routes

Because high-copy-number routes are frequently underdetermined, we next asked whether their bounded timing intervals retained the event times independently supported by determined routes. To do so, we checked whether underdetermined solution spaces were consistent with WGD and ccPG time points inferred from determined routes alone. In both illustrative tumors, the bounded solution spaces of underdetermined routes contained the event times inferred from determined routes. Specifically, exact timings obtained from determined routes for the (2, 0), (2, 1), and (3, 1) states fell within the underdetermined solution spaces for states ranging from (2, 2) to (7, 2) in the stomach tumor and up to (5, 0) in the esophageal tumor (**Figure 2a,b**, right). This concordance is possible because many underdetermined routes admit special-case solutions in which gains on distinct allelic branches occur simultaneously (**Supplementary Figure S4a**), as expected during WGD. Across the PCAWG cohort, these simultaneous-allele special-case solutions showed greater concordance with WGD time points inferred from determined routes than did randomly selected admissible solutions from the underdetermined solution spaces (**Supplementary Figure S4b**).

Finally, we benchmarked the accuracy of predictions for both determined and underdetermined solutions across copy number states using simulated cancer genomes. Our results showed similar error rates for determined and underdetermined solutions (**Supplementary Figure S3e**), indicating that copy-number-state complexity, rather than determinacy itself, was the principal driver of error (**Supplementary Figure S3f**). Higher-copy-number states partition molecular time into finer intervals and therefore yield narrower estimated solution ranges, making those ranges more likely to exclude the true timing in the presence of sampling error (**Supplementary Figure S5a**). Importantly, additional mutations reduced sampling error in the estimated constraints, but they cannot eliminate structural indeterminacy (**Supplementary Figure S5b**).

Together, the empirical concordance and simulation benchmarks show that Tau accurately estimates gain times for determined routes and provides informative bounds for underdetermined routes.

### Genome-wide timing distinguishes WGD from other punctuated multichromosomal gains

Extending the two illustrative examples, we applied Tau across PCAWG to identify punctuated multichromosomal gain events and distinguish WGDs from ccPGs. Gains linked to these two event classes jointly accounted for 78% of all clonal copy-number gains (**Figure 3a**), highlighting their central role in copy-number evolution. Given this substantial contribution, we first assessed Tau’s ability to distinguish and time the two event classes in simulated tumors (**Supplementary Figure S6a**). Tau accurately recovered both WGDs and ccPGs and their timing (**Figure 3b,c**), with high sensitivity and a low false-positive rate for events separated by at least 0.1 molecular-time units (**Supplementary Figure S6b,c**). We then characterized the prevalence, genomic extent, and composition of these events across PCAWG.

**Figure 3:**
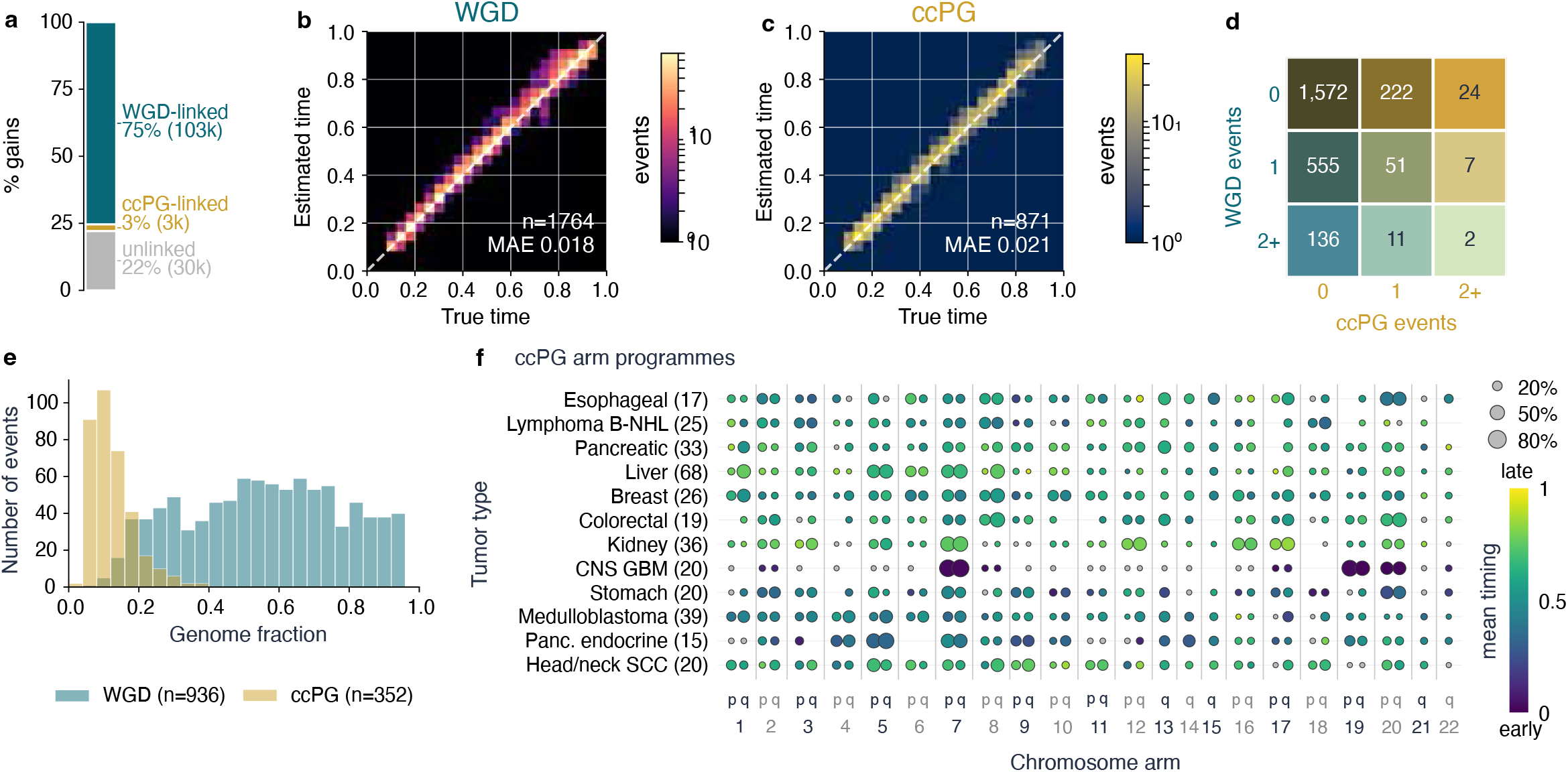
Genome-wide timing distinguishes WGD from cross-chromosomal punctuated gains (ccPGs). **a**, Partition of all clonal gains in PCAWG into WGD-linked, ccPG-linked and unlinked categories. **b**,**c**, Timing accuracy in simulated tumors: estimated against true event time for whole-genome duplications (WGDs, **b**) and cross-chromosomal punctuated gains (ccPGs, **c**). Color, number of events (log scale); MAE, median absolute error. **d**, Number of PCAWG samples by number of detected WGD and ccPG events. **e**, Fraction of the genome affected per event, for WGD and ccPG. **f**, Chromosome arms contributing to ccPGs, by tumor type. Dot size, percentage of that type’s ccPG events involving the arm; color, mean molecular time of those events.

At least one WGD was detected in 29.5% of PCAWG samples (**Figure 3d**). Individual WGDs affected a median of 55.5% of the genome (IQR, 37.0–73.4%) and involved substantial fractions of biallelic gain (**Supplementary Figure S6d,e**). By contrast, ccPGs were detected in 12.3% of samples, affected a median of 12.4% of the genome (IQR, 7.8–15.1%; **Figure 3e**), and predominantly involved gains affecting a single chromosomal copy (**Supplementary Figure S6d,e**). Approximately one-fifth of samples with either a WGD or ccPG harbored a second punctuated event (**Figure 3d**).

The chromosome-arm composition of ccPGs was strongly tumor-type-specific (**Figure 3f**). In glioblastoma, which had the highest ccPG prevalence, these events occurred particularly early and most commonly involved chromosome 7, where *EGFR* resides, frequently together with chromosomes 19 and 20 (**Figure 3f**; **Figure 4a,b**). In kidney cancers,ccPG prevalence exceeded WGD prevalence, and the events involved a distinct combination of chromosomes 7, 12, 16, and 17. Unlike the early events in glioblastoma, kidney ccPGs occurred across a broad range of later molecular times (**Supplementary Figure S7**). Thus, ccPGs constitute a distinct, predominantly monoallelic class of punctuated gain whose chromosomal targets and timing vary across tumor types.

**Figure 4:**
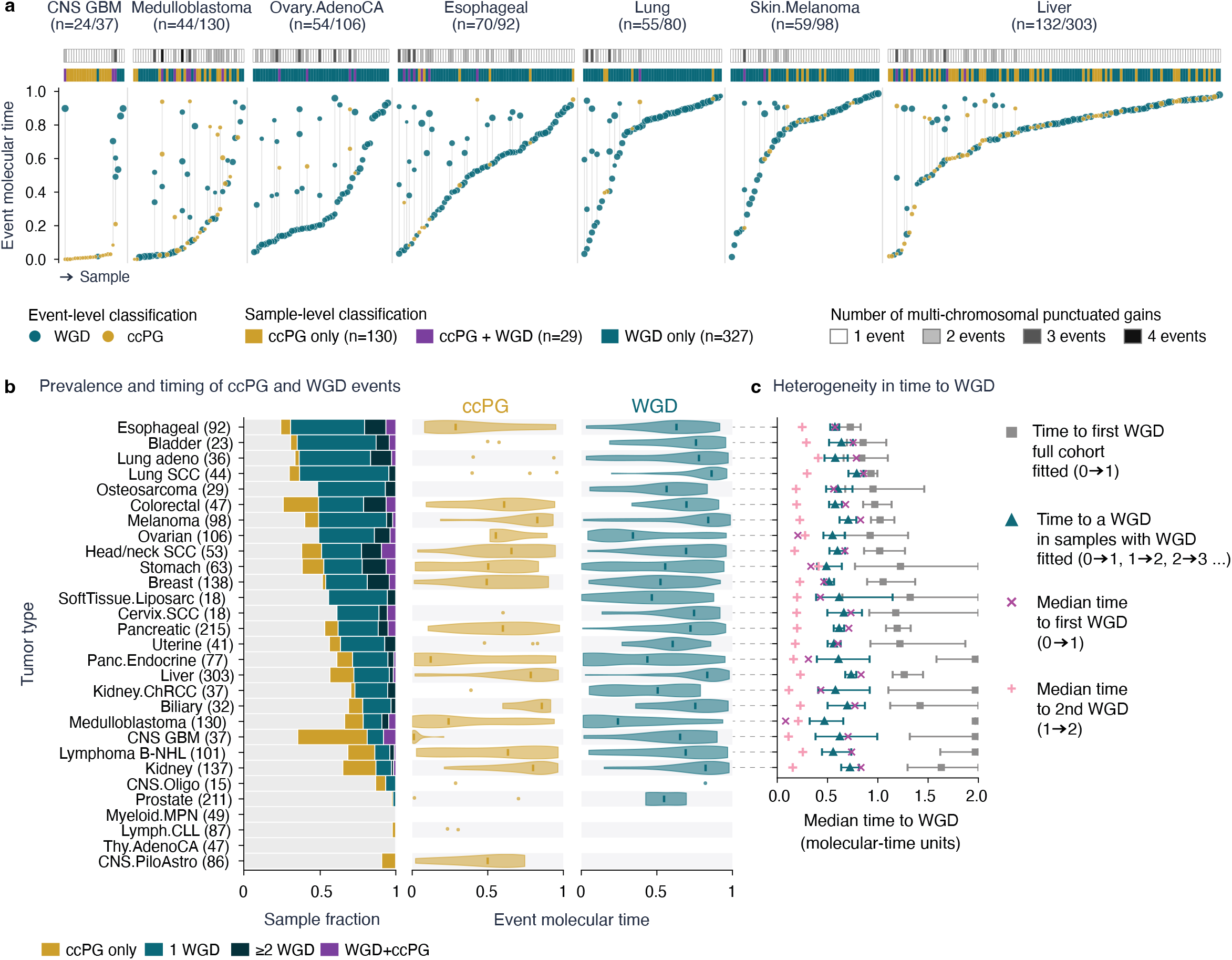
WGD and ccPG prevalence and WGD propensity are highly heterogeneous within and across tumor types. **a**, Per-sample event times in tumor types with frequent events; each column is a sample, ordered by event time, with WGD (blue) and ccPG (gold) events. Tracks above indicate event composition and number of events per sample. **b**, Prevalence of WGD and ccPG by tumor type (left) and the distribution of ccPG (center) and WGD (right) event times. **c**, Median molecular time to WGD by tumor type, comparing estimates from the full cohort with those restricted to samples that acquire a WGD, and the time to a first versus a second WGD. Only tumor types with at least three WGD events were included.

### Timing of multiple WGDs reveals heterogeneous genome-doubling dynamics

WGD prevalence varied markedly among tumor types, exceeding 60% in esophageal, bladder, and lung cancers but reaching only 2% in prostate cancer (**Figure 4b**). WGD dynamics also varied among tumors: some underwent multiple WGDs, whereas others underwent none (**Figure 4a**). Even among WGD-positive tumors, ranked distributions of first-WGD times showed pronounced changes in slope, including concave patterns in lung cancers, melanomas, and liver cancers and convex patterns in ovarian cancers and pancreatic endocrine tumors (**Figure 4a**; **Supplementary Figure S7**). These patterns suggested that WGD propensity may vary even among tumors of the same type, with genome-doubling-prone tumors forming a distinct subset rather than all tumors having an equal probability of WGD.

To quantify this heterogeneity, we modeled the waiting time to WGD while accounting for tumors in which no WGD was observed. Across the full cohort, the model-extrapolated median time to first WGD was 1.57 molecular-time units (95% CI, 1.45–1.69). This estimate extends beyond the observable 0–1 molecular-time interval, reflecting that fewer than half of all tumors acquired WGD during the reconstructable clonal lineage. By contrast, a transition model incorporating intervals to first and subsequent WGDs among WGD-positive tumors estimated a median waiting time of 0.59 molecular-time units (95% CI, 0.573–0.616). Although conditioning on an observed WGD contributes to this difference, the substantially shorter conditional estimate is consistent with a subset of tumors having an increased propensity for genome doubling.

We next asked whether this pattern was explained solely by differences among tumor types. We therefore repeated the comparison within each tumor type, contrasting the estimated time to first WGD across all tumors with the corresponding estimate restricted to WGD-positive tumors (**Figure 4c**). In tumor types with high WGD prevalence, including esophageal, bladder, and lung cancers, the estimates converged, consistent with a large proportion of tumors having a high propensity for genome doubling. The gap widened progressively in tumor types with lower WGD prevalence, suggesting that WGD propensity is confined to a smaller subset of tumors in these lineages. Moreover, among WGD-positive tumors, the median waiting time to the next WGD varied relatively little across tumor types (0.47–0.79 molecular-time units). Thus, with respect to WGD timing, tumors that undergo genome doubling across different cancer types resemble one another more closely than they resemble WGD-negative tumors of the same type.

We then tested whether the proposed WGD-prone state was associated with more rapid recurrence. Among tumors with multiple WGDs, the interval between the first and second WGD was generally shorter than the interval from tumor initiation to the first WGD, indicating an increased rate of genome doubling after the initial event (**Figure 4c**). Ovarian cancers and medulloblastomas were notable exceptions: their first WGDs occurred particularly early, but an early first WGD did not predict rapid acquisition of a second, and the interval between WGDs was, on average, longer than the time preceding the first. This pattern is more consistent with WGD acting as an early, tumor-type-specific selective event in these lineages than with a persistent propensity for repeated genome doubling.

By resolving multiple WGDs within individual tumors, Tau enabled the first systematic pan-cancer analysis of successive clonal WGDs preceding the MRCA. This analysis revealed marked heterogeneity in the timing and recurrence of genome doubling, distinguishing tumors with a persistent propensity for repeated WGD from lineages in which WGD appears to act as an early selective event.

### Timing high-copy-number amplifications reveals bursts enriched around WGD

Because Tau resolves the timing of every successive gain contributing to a high-copy-number state, it can reconstruct amplification trajectories that were previously only partially accessible, generally at lower copy numbers or through their earliest and latest events. To place these trajectories in a genome-wide context, we classified all clonal gains as WGD-linked, ccPG-linked, or unlinked to either event class and mapped their aggregated temporal and genomic distributions within each tumor type (**Figure 5a**; **Supplementary Figure S8**). All three gain classes contributed to the landscapes of highly amplified regions. Unlinked gains were distributed broadly across molecular time and chromosomes but retained tumor-type-specific concentrations at particular loci, including early gains on chromosome 3q in head and neck cancers and chromosome 7 in glioblastomas. These regions overlapped those targeted by ccPGs in the corresponding tumor types, suggesting that coordinated and isolated WGD-independent gains converge on common chromosomal targets. Whereas ccPGs were predominantly arm-level, unlinked gains spanned a wider genomic range, from whole-arm aneuploidies to focal amplifications affecting narrow genomic regions.

**Figure 5:**
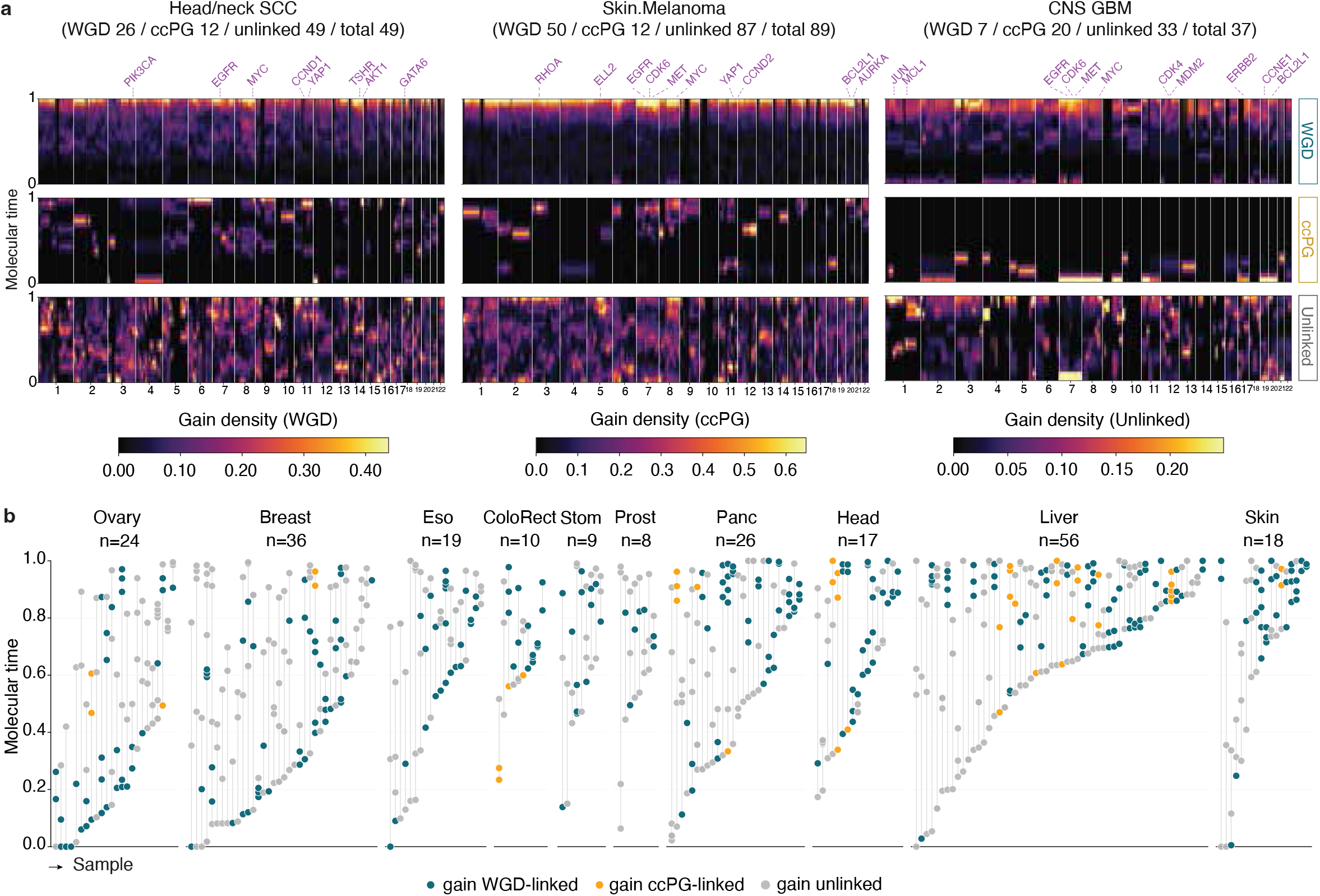
Temporal and genomic landscapes of WGD-linked, ccPG-linked and unlinked gains. **a**, Gain density across the genome (*x*) and molecular time (*y*) for three representative tumor types, partitioned into three layers: WGD-linked (within 0.20 molecular time of a detected WGD), ccPG-linked (within 0.10 of a ccPG and not already WGD-linked), and unlinked. Each layer is normalized by its own per-locus segment denominator, so color denotes the density of gains per contributing segment; scales are shared across tumor types within a layer but not between layers. Headers give the number of samples contributing to each layer. Selected oncogene loci are marked above. **b**, Sequence of gains at *MYC*. Each column is a sample; points give the molecular time of every gain at that locus, colored by whether the gain is WGD-linked (blue), ccPG-linked (gold) or unlinked (grey). Samples are ordered by the time of their first gain. Gains dispersed across long intervals indicate gradual amplification, whereas points stacked within a narrow interval indicate a burst.

Although WGD acts across the genome, its retained copy-number effect was not uniform. Part of this genome-wide heterogeneity reflects locus-specific copy-number histories before and after WGD. Gains acquired before WGD create a higher local copy number for genome doubling to amplify, whereas losses after WGD reduce the number of doubled copies that remain. Because WGD is expected to duplicate every existing chromosomal copy, we inferred post-WGD loss when an allele retained fewer copies than expected from doubling its pre-WGD state (**Supplementary Figure S9a**). The inferred losses were unevenly distributed across the genome, and recurrently affected regions varied among tumor types, thereby contributing to the observed WGD-linked landscapes (**Supplementary Figure S9b**). Nevertheless, WGD-linked gain density remained particularly heterogeneous at oncogenic loci, even after accounting for these considerations, motivating us to test whether WGD coincided with localized amplification beyond that expected from passive doubling alone.

Recurrently amplified high-copy-number regions frequently contained oncogenes. We reconstructed successive gains at loci including *MYC, PIK3CA*, and *EGFR* (**Figure 5b**; **Supplementary Figure S10**), where WGD-linked gains made a substantial contribution. The resulting trajectories followed two broad patterns: some tumors accumulated gains gradually across extended molecular-time intervals, whereas others acquired multiple gains within a narrow interval (**Figure 5b**). Although the term “burst” has previously been applied to punctuated gains shared across genomic segments, here we use it specifically to describe closely timed gains along a single evolutionary route. Some bursts occurred independently of WGD, either in WGD-negative tumors or at times separated from WGD, whereas many clustered around WGD time points. We operationally defined a burst as at least three consecutive gains occurring within 0.05 molecular-time units; trajectories not meeting this criterion were classified as gradual (**Figure 6a**; **Supplementary Figure S11**; Methods). Matched null simulations preserving copy-number states and mutation counts produced substantially fewer bursts than observed in PCAWG, indicating that the detected patterns were not explained by random spacing of successive gains (**Supplementary Figure S12a**).

**Figure 6:**
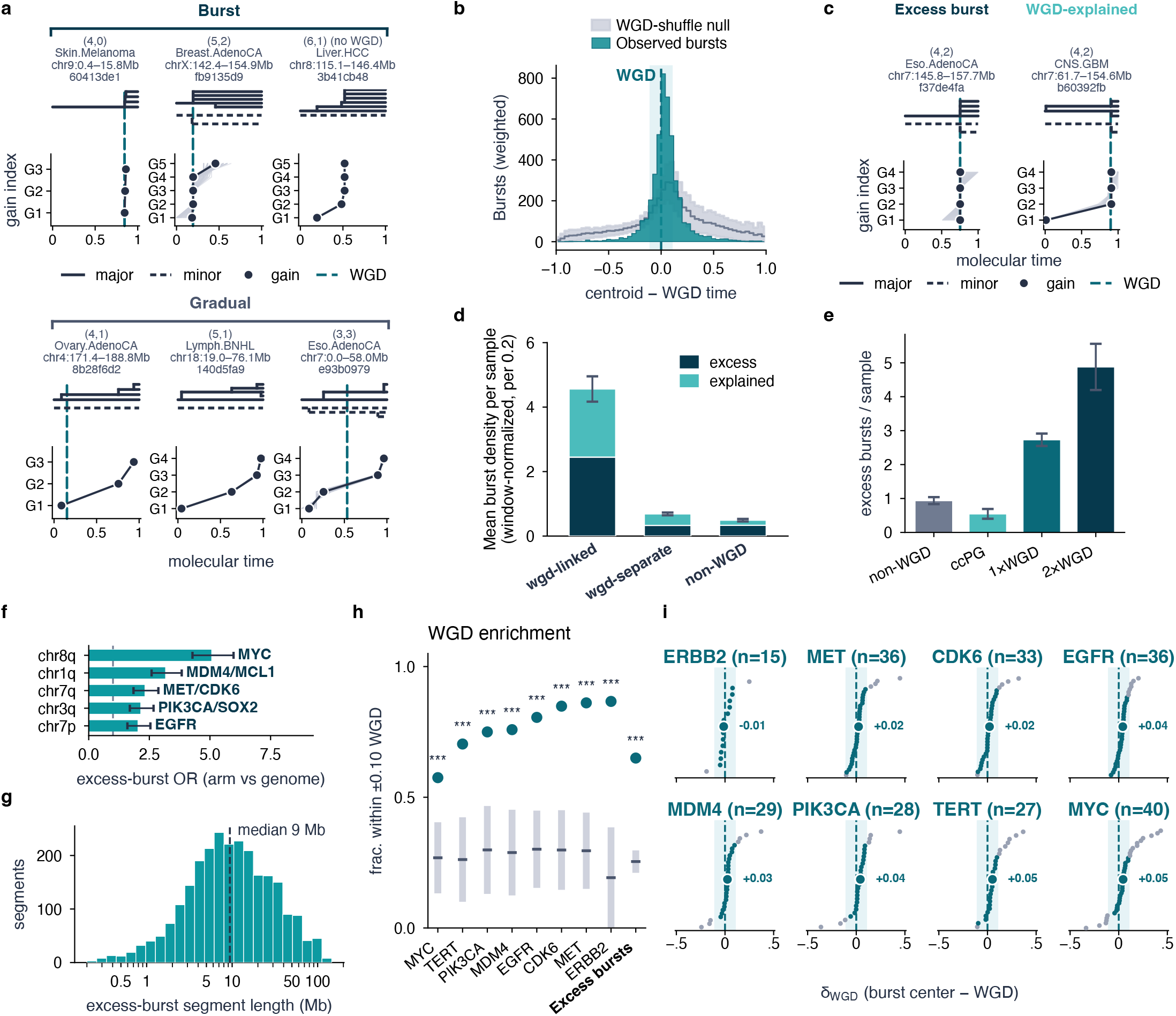
High-copy-number amplifications occur in bursts enriched around whole-genome duplication. **a**, Representative high-copy-number segments showing burst (top) and gradual (bottom) amplification. For each, the allele tree (major, solid; minor, dashed) is shown above the inferred time of each successive gain (gain index against molecular time); shading gives the range of solutions and the dashed line the sample’s WGD. **b**, Distribution of the time between a burst and the nearest WGD, for observed bursts and for a null generated by permuting WGD times between samples. **c**, Bursts containing more gains than the maximum attributable to a single genome doubling (excess, left), compared with bursts containing a number of gains compatible with doubling alone (WGD-explained, right). **d**, Mean burst density per sample, normalized per 0.2 units of molecular time, split into excess and WGD-explained bursts, for WGD-linked and WGD-separate windows in WGD samples and for samples without WGD. **e**, Excess bursts per sample by sample class. Error bars, s.e.m. **f**, Chromosome arms enriched for excess bursts relative to the genome average, with the oncogenes they contain marked. **g**, Length distribution of segments carrying excess bursts. **h**, Fraction of excess bursts falling within ±0.10 molecular time of a WGD, per oncogene and across all excess bursts, against a permutation null (grey). ^***^*P <* 0.001. **i**, Time from each burst center to the nearest WGD (*δ*_WGD_) per oncogene; labels give the median offset.

Strikingly, bursts were strongly enriched near WGD (**Figure 6b**). In a permutation null in which WGD times were shuffled among samples, bursts occurred near WGD substantially less often than in the observed data. Because WGD can duplicate gains already present at a locus, we calculated, for each final copy-number state, the maximum number of gains attributable to a single genome doubling and classified bursts containing additional gains as excess bursts (**Figure 6c**; Methods). Excess-burst density was 7.43-fold higher within a ±0.1 molecular-time window around WGD than in WGD-distal intervals from the same samples and 7.35-fold higher than in samples without WGD (**Figure 6d**). Excess-burst counts per sample also increased with the number of detected WGDs (**Figure 6e**). These results indicate that WGD marks a temporal window in which localized amplification beyond passive genome doubling occurs substantially more frequently.

Chromosome-arm enrichment analysis identified recurrent excess-burst hotspots, the most prominent of which overlapped established oncogenes (**Figure 6f**). Overall, segments containing excess bursts had a median length of 9.4 Mb (**Figure 6g**). At hotspot oncogenic loci, permutation analysis confirmed that the temporal alignment between bursts and WGD exceeded that expected by chance (**Figure 6h**). Moreover, several bursts fell just outside the prede-fined WGD window while remaining close to WGD time (**Figure 6i**), suggesting that the estimated fold enrichment may be conservative. Together, these results show that WGD frequently coincides with a window of localized, burst-like oncogene amplification that extends beyond the copy-number increase produced by genome doubling alone.

### Successive gain timing resolves pre-MRCA mutational-process dynamics

The same temporal resolution that allowed us to order multiple WGDs and reconstruct the successive gains leading to high-copy-number oncogene amplifications also enabled us to examine another dimension of tumor evolution: changes in mutational-process activity. Thus far in our analysis, clock-like SBS1/SBS5 mutations supplied the molecular timescale used to reconstruct successive gains; we next reversed this relationship and used the timed gains to partition the pre-MRCA clonal lineage into ordered intervals, allowing changes in other mutational processes to be traced.

Mutational-process activities can shift markedly during tumor evolution. Tobacco-associated mutagenesis in lung cancer is often concentrated early, whereas APOBEC-associated mutagenesis frequently becomes more prominent later and has been implicated in treatment resistance [3, 11, 30–32]. Unlike WGD-based early-versus-late classifications [11], Tau uses each successive gain as a temporal landmark, enabling signature timing at finer resolution and in tumors with or without WGD. Resolving gains within high-copy-number states increases the number of temporal landmarks, while route clustering by pooling segments with shared amplification histories increases the number of informative mutations within each interval. Tau then quantifies the accumulation of mutations attributed to each signature relative to the SBS1/SBS5 clock-like baseline within these gain-defined intervals.

Our approach uses process-weighted interval estimates directly, rather than first assigning mutations to temporal intervals based on multiplicity and then performing signature analysis within those groups, because a given multiplicity can correspond to multiple intervals. To see how, consider, for instance, the simple (2, 1) copy-number state. A mutation acquired on the minor allele before gain of the major-allele and a mutation acquired on one of the major-allele copies after that gain can both have a final multiplicity of one (**Supplementary Figure S13a**). Mutations with multiplicity one are therefore enriched for, but not unique to, the post-gain interval. Rather than assigning each mutation deterministically to one interval, Tau accounts for this ambiguity using the complete route structure and observed multiplicity distribution. For each mutational process, Tau weights every mutation by its probability of having been generated by that signature and substitutes the resulting signature-weighted multiplicity counts into the same route-specific system of linear equations used to infer SBS1/SBS5 molecular-time (Methods). This produces a process-specific estimate for each time interval, which Tau compares with the corresponding SBS1/SBS5 interval. Higher signature-to-clock time interval ratios, temporal dilations, indicate an increase in relative process activity, whereas lower ratios, temporal contractions, indicate a decrease in relative process activity in this interval (**Supplementary Figure S13c**). Tracking this ratio across successive gain-defined intervals reconstructs temporal changes in relative mutational-process activity along the pre-MRCA clonal lineage.

We illustrate this approach using the esophageal adenocarcinoma example introduced above (**Figure 2b**). Mapping SBS17 activity onto the same segment-level timing structure (**Figure 7a**, left) and subsequently onto the clustered evolutionary routes (**Figure 7a**, right) revealed a consistent temporal pattern across segments and routes, with SBS17 enriched during early intervals and progressively depleted at later molecular times. To systematically identify such changes, Tau fits a stepwise-change model to route-level relative signature rates and uses the Bayesian information criterion to determine the number of activity transitions supported within each sample (**Supplementary Figure S13d,e**). Applying this framework across PCAWG revealed recurrent, but tumor-type-specific, temporal trajectories of mutational processes. Most esophageal adenocarcinomas showed early enrichment followed by declining SBS17 activity, whereas in stomach cancers SBS17 was prominently observed as a late process (**Figure 7b**, left). SBS17 is a dominant mutational process in esophageal adenocarcinoma linked to oxidation of the nucleotide pool [33, 34], and it has been previously reported to be active early in tumor development, including in Barrett’s esophagus, a precursor condition associated with increased esophageal-cancer risk [33, 35, 36]. The SBS17 activity drop we observe over time is therefore consistent with previous observations spanning premalignant and subclonal disease. These trends were not uniform across tumors, however, revealing substantial inter-tumor heterogeneity in the temporal deployment of the same mutational processes. A subset of esophageal adenocarcinomas and most stomach adenocarcinomas displayed the opposite SBS17 trajectory, with activity increasing toward the MRCA. Such cases suggest that SBS17 activity is not invariably confined to early tumor initiation and may instead remain active, or become more prominent, during later clonal expansions preceding the MRCA.

**Figure 7:**
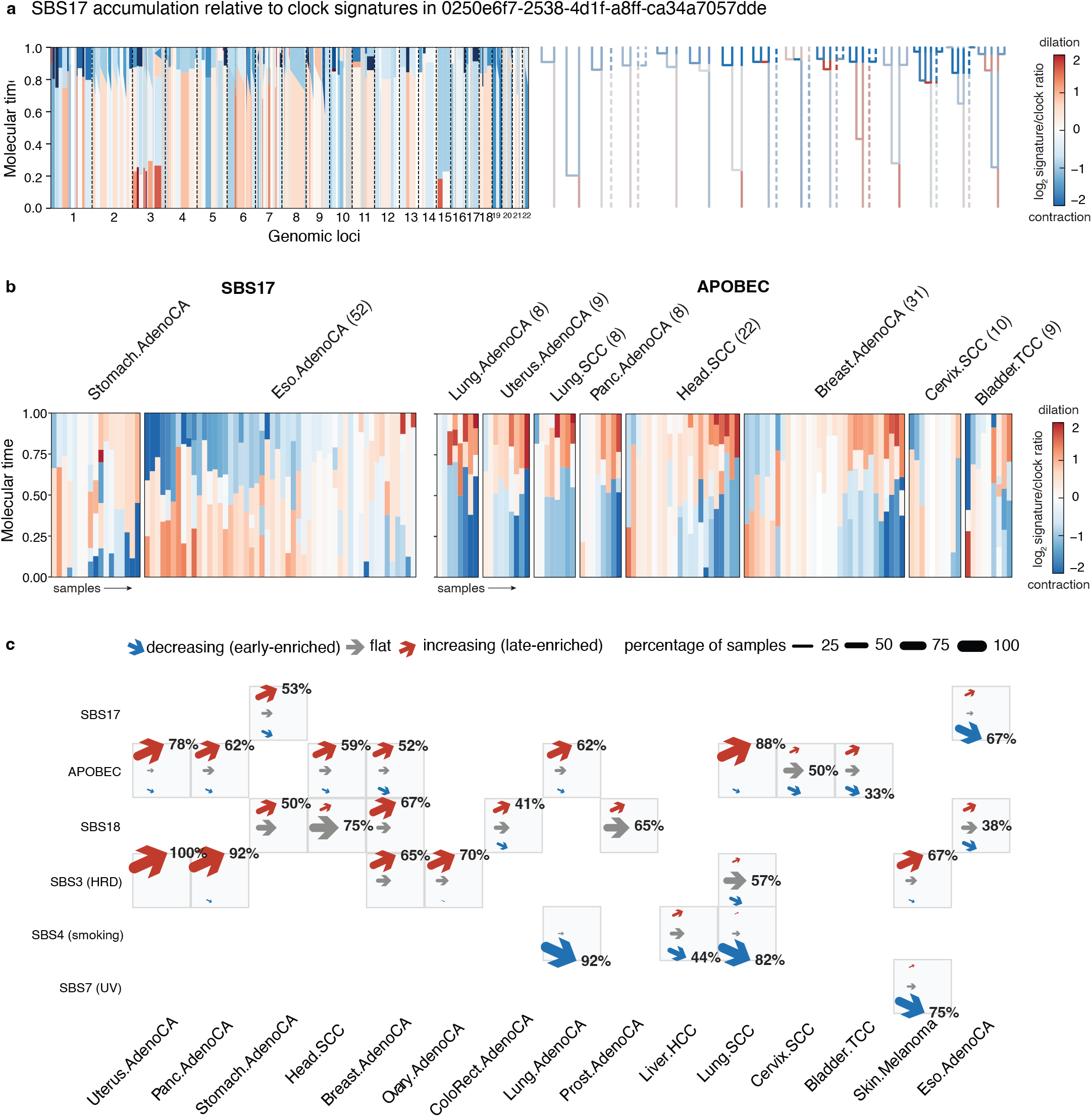
Mutational-process activity resolved across clonal history. **a**, SBS17 activity relative to the SBS1/SBS5 clock across the genome (*x*) and molecular time (*y*) in the esophageal adenocarcinoma of **Figure 2b**. Color, log_2_ of the signature-to-clock ratio per interval; red indicates intervals in which SBS17 accumulated faster than the clock and blue slower. SBS17 is enriched early and depleted later. On the right are trees representing the amplification routes of pooled-segment clusters, colored by SBS17 activity during different intervals. **b**, The same ratio as in panel (a) for SBS17 and APOBEC across all PCAWG tumor types. Samples are ordered from early to late enrichment from left to right, with molecular time on the y-axis. **c**, Direction of signature-activity change across successive pre-MRCA intervals, by signature and tumor type. Arrows denote the proportion of samples whose trajectory was classified as decreasing (blue), flat (grey) or increasing (red); arrow size is proportional to that fraction and the label gives the dominant class. Cells are shown only for tumor type–signature combinations with at least eight classified samples.

Consistent with expectations, APOBEC-associated SBS2 and SBS13 generally showed the opposite trajectory, with increasing relative activity toward later molecular-time intervals (**Figure 7b**, right). Similarly, although late enrichment of APOBEC-associated mutagenesis was common across tumor types, this pattern was more heterogeneous in tumor types such as breast, bladder, cervical, and head and neck cancers, where a subset of tumors showed substantial APOBEC activity during earlier evolutionary intervals. Particularly in cervical and bladder cancers, this early enrichment is consistent with a potential role for APOBEC mutagenesis in tumor initiation, supported by evidence of virus-induced APOBEC activation and APOBEC-mediated DNA damage in nonmalignant cells [37–39].

We expanded the analysis across tumor types and classified sample-specific signature trajectories as increasing, decreasing, or flat. Several mutational processes associated with carcinogenic exposures or early tumor development—including tobacco-associated SBS4 in lung cancer, ultraviolet-associated SBS7 in melanoma, and SBS17 in esophageal adenocarcinoma—showed their highest activity in the earliest pre-MRCA intervals (**Figure 7c**), therefore decreasing over time. Conversely, processes such as homologous recombination deficiency, which typically arise after a sequence of genomic alterations, including *TP53* mutations and loss of heterozygosity of *BRCA1* or *BRCA2* in carriers of germline mutations in these genes, tended to emerge toward the end of the pre-MRCA time period, thus increasing over time. This timing is consistent with these processes being a consequence of preceding genomic events rather than an initiating trigger. Together, these results demonstrate that copy-number amplification histories provide temporal landmarks for resolving changes in mutational processes within the previously compressed pre-MRCA period, revealing both recurrent tumor-type-specific trajectories and substantial heterogeneity in when individual mutational processes become active during cancer evolution.

## Discussion

Copy-number timing methods have established that mutation multiplicities can reconstruct gain histories along the clonal lineage of a tumor [11, 22–24]. Tau generalizes this principle by enumerating allele-specific gain routes and deriving exact or bounded analytical timing solutions for successive gains. This formulation retains non-identifiable histories rather than forcing constraints based on preconceived assumptions, while precomputed solutions make exhaustive route evaluation computationally tractable. Pooling compatible segments and clustering their gain times connects segment-level histories to genome-wide events, which allows WGD and other punctuated gains to be identified after unrestricted route inference and enables multiple such events to be resolved within individual tumors. We compiled the resulting sample-level reconstructions into a browsable resource for exploring amplification histories across the widely studied PCAWG cohort.

The ability to resolve multiple punctuated events exposed substantial heterogeneity in genome-doubling dynamics. The expected waiting time to WGD across the full cohort was considerably longer than that among WGD-positive tumors, whereas the interval between successive WGDs was generally shorter than the interval preceding the first. These patterns suggest that a subset of tumors enters, or is selected for, a WGD-prone state in which repeated doubling becomes more likely. WGD-positive tumors across cancer types also resembled one another in their timing more closely than they resembled WGD-negative tumors of the same type. Supporting a view of WGD as a dynamic rather than one-time process, recent single-cell analysis of high-grade serous ovarian cancer identified early fixation, multiple parallel events, and late subclonal genome doubling [40]. Whereas Tau resolves successive clonal WGDs from bulk genomes, single-cell analysis can additionally capture parallel and subclonal doubling events. Ovarian cancers and medulloblastomas were exceptions to the general association between early first WGD and rapid recurrence: their first WGDs occurred particularly early but did not predict accelerated acquisition of a second clonal WGD. In ovarian cancer, this may indicate that subsequent doublings arise later and independently across subclonal branches rather than as successive events shared by the tumor population.

Beyond WGD, Tau identified recurrent ccPGs, which were more restricted, predominantly monoallelic, and composed of tumor-type-specific combinations of chromosome arms. Together, WGD and ccPG events accounted for 78% of copy-number gains across PCAWG, indicating that punctuated amplification extends beyond genome doubling. The predominantly monoallelic, arm-level structure of ccPGs is compatible with transient chromosome mis-segregation, coordinated rearrangement, or selection for particular combinations of arm-level gains. Their potential biological importance was especially apparent in glioblastoma, where early ccPGs recurrently involved chromosome 7, often together with chromosomes 19 and 20. Despite their different scales, WGD-linked, ccPG-linked, and unlinked gains converged on overlapping tumor-type-specific regions containing established oncogenes.

Within highly amplified regions, Tau resolved successive gains and showed that copy-number expansion frequently occurred through temporally concentrated bursts rather than gradual accumulation. These bursts were more common than expected under matched random-amplification models and were strongly enriched near WGD. By comparing each burst’s gain count with the maximum gain count a WGD event could produce, we distinguished gains attributable to genome doubling from excess gains that required additional amplification events. Many WGD-proximal bursts contained such excess gains, and the excess-burst burden increased with the number of WGDs within a tumor. Moreover, recurrent excess-burst hotspots overlapped established oncogenes, suggesting that these periods of accelerated amplification preferentially increase dosage at selected cancer-associated loci. WGD could create a transient state permissive for localized amplification through increased genomic instability or greater tolerance of dosage imbalance. Alternatively, WGD and localized bursts may be parallel consequences of a shared unstable state, with selection retaining advantageous oncogene amplifications. Although their temporal association does not establish causality, these findings show that WGD frequently coincides with localized oncogene amplification beyond passive doubling of the pre-existing genome.

Successive gain times also partitioned the clonal lineage into ordered intervals, allowing changes in mutational-process activity before the MRCA to be resolved. Processes associated with early carcinogenic exposure or tumor development—including SBS4 in lung cancer, SBS7 in melanoma, and SBS17 in esophageal adenocarcinoma—generally declined across successive intervals. In contrast, APOBEC-associated SBS2 and SBS13 more often increased toward later pre-MRCA evolution, consistent with their reported contribution to later tumor diversification [3, 11, 32, 41]. These patterns were not universal, however, revealing substantial intertumor heterogeneity in when the same process became active. Tau therefore extends broad early-versus-late classifications by resolving multiple intervals within the shared clonal lineage. This information is complementary to TrackSig and related cancer-cell-fraction-based approaches [42, 43], which resolve subclonal trajectories but cannot readily order mutations that remain clonal at sampling.

Several limitations define the current scope of Tau and motivate future extensions. Timing accuracy depends on reliable allele-specific copy-number, purity, and SNV estimates and on sufficient clock-like mutations. Greater sequencing depth can reduce sampling error but cannot resolve structurally non-identifiable histories. Where available, orthogonal data from long-read, single-cell, or multiregion sequencing could help constrain balanced high-copy-number states and distinguish synchronous special-case solutions that are supported but not uniquely determined by cross-segment evidence. Extending Tau to subclonal mutations and multisample or longitudinal cohorts with resolved phylogenies could enable gains to be timed along individual evolutionary branches rather than only along the shared lineage leading to the MRCA. Calibration using longitudinal observations or well-characterized age-associated mutation rates could similarly help relate mutation-based molecular time to chronological time. Finally, integrating timed gain trajectories with amplification architecture and experimental models could clarify the physical mechanisms underlying ccPGs and localized bursts and test whether WGD directly facilitates oncogene amplification or whether both arise from a shared state of genomic instability. These extensions require substantial methodological development and, in some cases, experimental validation; they therefore fall beyond the scope of the present study and represent directions for future work.

Together, these findings show that amplification histories comprise distinct and often punctuated modes whose timing and recurrence vary among tumors. By resolving successive gains without prespecifying WGD, Tau reconstructs high-copy-number evolution and reveals coordinated ccPGs, heterogeneous genome-doubling dynamics, WGD-associated oncogene-amplification bursts, and changes in mutational processes within the previously compressed pre-MRCA lineage.

## Methods

### Enumeration and analytical solution of all evolutionary routes to all copy-number states

Methods for timing copy-number amplifications in bulk whole genome sequencing (WGS) generally rely on the same simple intuition - that clonal somatic SNV mutations that occurred before a copy number amplification will be present on more copies than those that occurred after the amplification, hence the multiplicity (the number of copies on which a SNV is present) of a SNV acquired before an amplification will be higher. This results in a proportionally higher VAF. This can be generalized to any copy number state (CN_maj_, CN_min_), such that we can formulate as a system of linear equations (or equivalent matrix) any route to any copy-number state by relating the frequencies of different SNV multiplicities {*N*_*i*_} to the timing of gains {*t*_*i*_}. To generate all route matrices, amplification routes are broken down into nested routes of smaller states, enabling us to generate matrix representations of all possible routes to any copy number state, constructed recursively as products of allele-specific branching and propagation matrices (**Supplementary Figure S1b**). To solve these enumerated routes, we used SageMath [25] to pre-solve analytically all routes for {*t*_*i*_} up to a copy-number state of (7, 5). Hence, timing any specific segment requires only the minimal computation of substituting the segment’s numerical multiplicity counts into the analytical solution from the existing database. In addition to solving the system of linear equations, we included a set of constraints on our *t* variables (*t*_*i*_ ≥ 0), which mean the output solution not only includes equalities but also constraints on {*N*_*i*_} and {*t*_*i*_}, which reduce the solution space, and enable further inferences on the tumor’s evolution.

### Processing of SNVs to calculate SNV multiplicity counts

Tau’s analytical solutions rely on input of accurate estimations of {*N*_*i*_}, the set of SNV multiplicity counts per segment – in other words, how many SNVs are present on one segment copy, two segment copies, and so on. A SNV’s multiplicity will determine its VAF. Hence, per SNV, to estimate multiplicity, we must take into account the number of ALT reads (*x*), tumor purity (*ρ*) and read depth (*n*) and total copy-number CN_tot_ = CN_maj_ + CN_min_. This requires pre-existing copy-number calls. For PCAWG tumor samples, we used the PCAWG consensus CNA tumor purity and calls. We model the likelihood of VAF of SNV *i* having multiplicity *j* as follows:

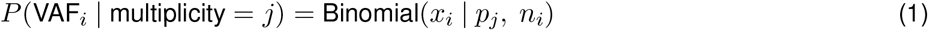

Where

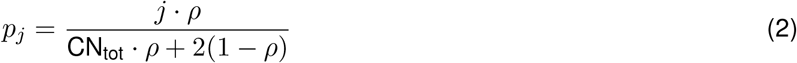

We estimate the distribution *P*(*m*_*i*_ = *j*) = *π*_*j*_ in order to generate the counts {*N*_*j*_} for Tau. The counts correspond to the expected number of SNVs of each multiplicity, based on our *π*_*j*_ and the total number of SNVs in the segment. Tau provides an optional heuristic filter to remove likely subclonal SNVs based on cancer cell fraction. We also provide the option to specify a custom list of subclonal SNVs to be removed before estimation. Then, following previous methods like MutationTimeR [11], we use Expectation-Maximization to estimate multiplicity counts {*N*_*i*_}. Optionally, one can skip multiplicity estimation entirely and specify custom multiplicity counts.

### Power-to-detect correction

Low-VAF mutations are less likely to be reported by the variant caller, which biases estimated multiplicity counts toward higher multiplicities. We correct for this by conditioning the per-SNV likelihood on passing the caller’s detection rule. Let *x*_*i*_ and *n*_*i*_ be the alternate read count and total depth of SNV *i*. For a segment with total copy number *CN*_tot_ = *CN*_maj_ + *CN*_min_ and purity *ρ*, the expected alternate fraction under multiplicity *j* is

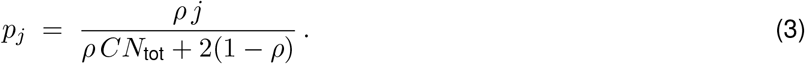

Under this model *X*_*i*_ ~ Binomial(*n*_*i*_, *p*_*j*_). If the caller requires at least *k*_*i*_ alternate reads for SNV *i* (by default *k*_*i*_ = 3), we use the truncated (conditional) binomial likelihood

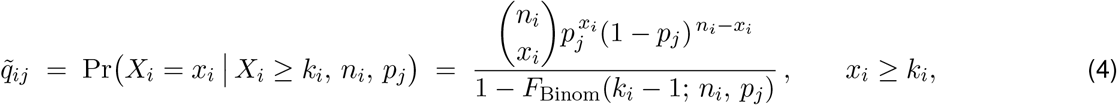

and 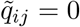. Here *F*_Binom_ is the binomial CDF.

Writing *q*_*ij*_ = Pr(*X*_*i*_ = *x*_*i*_ | *n*_*i*_, *p*_*j*_) for the uncorrected binomial likelihood used above, the posterior over multiplicities replaces *q*_*ij*_ with its power-corrected counterpart 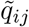:

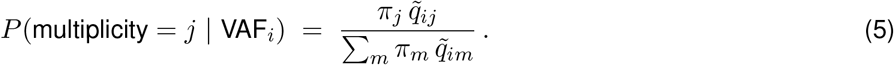

Conditioning on detection in this way removes the preferential loss of low-VAF mutations and yields power-corrected estimates of the segment-level multiplicity counts {*N*_*j*_}.

### Timing gains in individual segments

Once multiplicity counts have been estimated, they are substituted into the appropriate copy-number solution set to estimate timing results for all the segment’s amplifications. As discussed above, each analytical solution is accompanied by a set of constraints. Because there are multiple routes to the same copy-number, each route will have its own analytical solution with a unique set of constraints. Constraints relying purely on {*N*_*i*_} delineate which multiplicity frequency distributions are consistent with such a route and which are inconsistent. Such constraints therefore help to distinguish between different evolutionary routes to the same copy-number state. These constraints on {*N*_*i*_} can be applied to eliminate large fractions of routes without sampling-based inference. For example, a genomic segment of copy number state (4,2) can be achieved in multiple ways starting from a diploid state of (1,1). Whether the minor or major allele is amplified first, for instance, changes the observed multiplicity frequency space, and gives rise to a different set of equations, using {*N*_*i*_} (the number of SNVs of multiplicity *i*) to calculate {*t*_*j*_} (the set of time intervals between gains).

In addition, a distinct set of constraints, those involving {*t*_*i*_}, help narrow the solution space further. Depending on the rank of the route matrix and the copy-number state, the resulting solution will be exact or underdetermined. Routes to simpler copy-number states, such as (2, 1), have exact solutions. Routes to more complex copy-number states, such as (4, 2), will have one or more free variables, meaning there are many possible timings corresponding to a given SNV multiplicity frequency distribution, complicating the interpretation of results. Previous timing approaches have therefore often sidestepped these more complex cases. Tau, however, addresses these complex cases directly by harnessing the fact that although solutions are underdetermined, each *t*_*i*_ is still bounded. Knowing the full space of {*t*_*i*_} therefore still provides information on the timing landscape. To calculate this full timing space, we solve each system of linear equations with added constraints, namely the specification that each *t*_*i*_ ≥ 0. This produces an output solution set of both equalities (each *t*_*i*_ as a function of other variables) and inequalities (bounds on {*N*_*i*_} and {*t*_*i*_}).

This renders the timing of more complex copy-number states more tractable, as, for each possible route, we now have constrained ranges of timing determined by the observed SNV multiplicity counts. For instance, in the example case of a segment of copy-number (4, 2), there are one or two free variables, depending on the route, creating a constrained range of times for each amplification (**Figure 1d**, bottom). These constraints therefore both limit the space of feasible evolutionary routes, and provide upper and lower bounds to underdetermined timing results. Taking this into account, we can generate tree representations of the segment’s evolutionary history consistent with the data.

### Generating special-case solutions for underdetermined routes

For WGD samples, it is highly likely that there will be simultaneous gains. Therefore, for underdetermined timing solutions in samples with inferred WGD times, for visualization of routes and the concordance analysis (**Supplementary Figure S4**), we enforce a special-case solution which simply maximizes the simultaneity of gain times.

### Timing different mutational processes

#### Weighting SNVs by signature likelihoods and sample exposure

In order to estimate the multiplicity distributions of a specific mutational process, Tau weights SNVs by their probability of being generated by a given signature. Therefore, every SNV *i* is assigned a weight *w*_*ij*_, which corresponds to the probability of that SNV coming from mutational process *j*. This is achieved by taking signature exposures estimated by MuSiCal [44] and per-signature 96-channel trinucleotide mutational spectra from the COSMIC catalog [45] to calculate the probability of each SNV (SNV_*i*_) being generated by each signature (SBS_*j*_):

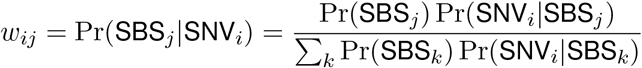

Pr(SNV_*i*_|SBS_*j*_) is taken directly from observing the likelihood of SNV_*i*_’s trinucleotide context as part of the signature SBS_*j*_ in the COSMIC mutational signature catalog. Pr(SBS_*j*_) is calculated by normalizing the exposures of all signatures detected in the sample from MuSiCal.

These probabilities are then used as weights in the EM estimation of multiplicity counts based on the signatures specified. If a mutational process is represented by multiple signatures, their respective weights are summed. By default, Tau uses both SBS1 and SBS5 to time segments, so as to estimate a clock-like multiplicity distribution and hence clock-like molecular time intervals. One can optionally include all mutations, or any chosen subset of signatures. This weighting enables the activity of different mutational processes to be analyzed during a tumor’s clonal evolution.

#### Comparing non-clock-like mutational process activity to clock-like activity

In order to investigate when a mutational process was active during a tumor’s development, Tau uses comparison of clock-like timing to the timing of some specific mutational process (**Supplementary Figure S13**). This is achieved by comparing, for a given interval in a given segment, the SBS1+SBS5-weighted interval *t*_clock_ and the mutational process-weighted *t*_sig_. The ratio 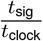 gives an estimate, during the interval spanned by *t*_clock_, for how active some mutational process was relative to clock-like time (**Supplementary Figure S13c**). If *t*_clock_ and *t*_sig_ have been normalized such that all *t*_clock_ and *t*_sig_ intervals within a segment respectively sum to 1, then the ratio only provides a measure of the relative activity of the process over time. If *t*_clock_ and *t*_sig_ are un-normalized, then it also provides a measure of how much more active the process was (i.e. the mutation rate) relative to the clock-like process during that interval. Un-normalized time interval values can be interpreted as an estimate of the number of mutations accumulated along one branch of the tumor’s evolutionary tree during that interval. For example, an interval of *t*_clock_ = 5 would imply 5 clock-like mutations accumulated per branch of the tree during that interval. Tau uses the un-normalized versions in order to preserve information about the absolute activity of a mutational process relative to the clock-like mutation rate.

Considering that single segments are noisy, Tau uses the pooled-segment clusters from clock-like timing in order to estimate mutational process activity over time. The corresponding segments within the clock-like cluster are then pooled and re-timed using the mutational process’s weights. Then the ratios are calculated for the mutational process-specific timing versus the clock-like timing per cluster. Different clusters will produce intervals with different start and end points, creating a series of overlapping intervals in molecular time. Each interval averages the true activity over that period. Tau combines this information across intervals to model the mutational process activity over time (**Supplementary Figure S13d,e**).

First, Tau sums up, over all clusters, all clock-like-weighted time intervals and all mutational process-weighted time intervals respectively. The ratio of these,

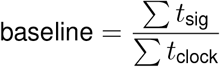

provides a measure of how active the mutational process of interest is compared to clock-like mutations averaged over all clusters and time intervals in the sample. Tau also calculates the log fold change per time interval *i*,

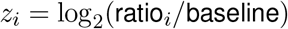

where ratio_*i*_ is the ratio of *t*_sig_ to *t*_clock_ in some time interval *i. z*_*i*_ is hence the interval’s mutational process activity relative to the average (baseline) over all clusters and time intervals. The final step remaining then is to infer the activity of the mutational process over time from these *z*_*i*_ values spanning overlapping intervals.

To achieve this, Tau fits a step function over molecular time. Molecular time is partitioned into *K* states. In state *k*, the mutational process is assumed to have constant activity *a*_*k*_. A fraction *f*_*ik*_ of interval *i* falls within state *k*. Then,

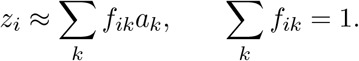

The activities *a*_*k*_ are estimated by weighted least squares with each interval weighted by the number of mutations attributed to the process within it, such that well-powered intervals contribute more to the fit. Starting from a single state, Tau adds a new state, with a changepoint that most reduces the weighted residual sum of squares (RSS). At each step, Tau calculates the Bayesian Information Criterion (BIC),

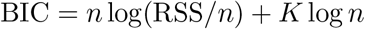

where *n* is the total number of intervals included in the fit. When the BIC no longer improves, the process stops adding new states, producing the lowest number of states that the data support to explain the mutational process activity over time.

In order to avoid overfitting the data, Tau enforces key constraints. No state may be narrower than 0.05 units of molecular time (normalized to 1), and intervals are only used if they contain at least 5 clock-like mutations and span between 0.05 and 0.75 units of molecular time. In addition, a state’s activity *a*_*k*_ must lie within the range of *z*_*i*_ observed during the state’s window of molecular time, restricted to what 10 or more mutations support. A sample is fitted only if its clock-like intervals sum up to at least 20 mutations across at least 5 intervals.

Per state, Tau reports the bounds on activity per state, log_2_ fold change with standard error, and the absolute activity per state, equal to the baseline × 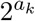.

#### Uncertainty of state estimates and classification of temporal trends

Writing **B** for the *n* × *K* matrix with entries *f*_*ik*_, **W** for the diagonal matrix of interval weights, a for the vector of state activities and **z** for the vector of *z*_*i*_, the weighted least-squares estimate is â = (**B**^T^**WB**)^*−*1^**B**^T^**Wz**. For the selected *K*-state model, residual variance was estimated as

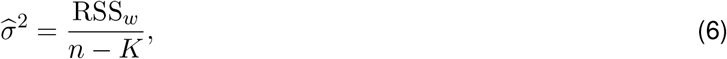

and the covariance matrix of the state estimates was

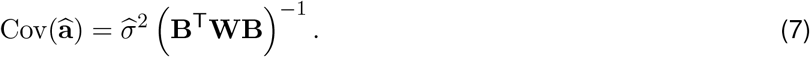

State-specific standard errors were obtained from the square roots of the diagonal elements. The uncertainty bands used in the per-sample profile plots (**Supplementary Figure S13d**,**e**) correspond to *âa*_*k*_ ± 1 standard error.

To summarize temporal direction across the cohort, we fitted a linear trend through the BIC-selected states for each sample and process. State *k* was represented by its midpoint *x*_*k*_ = (*l*_*k*_ + *h*_*k*_)*/*2 and fitted activity *â*_*k*_, and was weighted by 1*/* max(SE(*â*_*k*_), 0.001)^2^. Let *β*_1_ 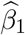 and 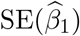 denote the resulting slope and its standard error. We classified a trajectory as increasing when 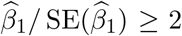, decreasing when this statistic was ≤ −2, and flat otherwise. Samples for which BIC selected a single state were classified as flat by construction. Tumor-type–signature combinations represented by fewer than eight classified samples were omitted from the pan-cancer directional summary. For each retained combination, arrow length and thickness represent the fraction of samples in each of the increasing, decreasing, and flat categories.

### Route selection

#### Different matrices encode different routes to the same copy-number state

A given copy-number state can be reached from an initial (1,1) state in several different ways. For example, the state (3,2) can be reached by major allele amplification first, and then a minor allele, e.g.

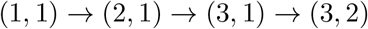

or it could be reached with a minor allele amplification first, e.g.

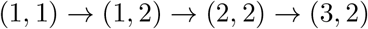

These different amplification orders (what we call “routes”) can give rise to different multiplicity states at different time intervals. To illustrate this, take the example above. Any mutation gained before the first major allele amplification will be observed ultimately with a multiplicity of 3. Therefore, the duration of time before the major allele amplification will determine the proportion of multiplicity 3 mutations. Hence, the major allele being amplified first implies the length of the first time interval (*t*_1_) is solely responsible for the proportion of multiplicity 3 mutations. Whereas, if the major allele is amplified after the minor allele, then the proportion of multiplicity 3 will depend on both the length of *t*_1_ and *t*_2_. These create two unique matrices for the different routes to (3, 2):

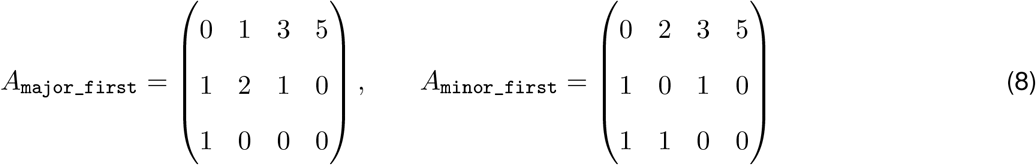

Each matrix, when solving for 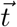 (time intervals between gains) based on observed 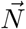 (multiplicity distribution), such that 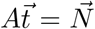, will give different solutions for 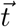. In addition, because of the imposed constraints 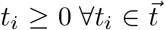, only a certain feasible region of the multiplicity space will give valid solutions, delineated by linear constraints calculated by SageMath [25].

#### Projection of observed multiplicities into the feasible region of a route

As explained above, Tau estimates 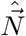 from SNV VAF distributions. This value is noisy, and so 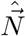, even if ultimately from some given route, may not satisfy all that route’s linear constraints. As the number of mutations increases, the size of violations decrease. We see similar distributions in simulations, suggesting that these constraint violations are largely a pattern of noise. In order to deal with this, we first of all normalize 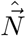 such that 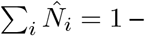 in effect, rescaling onto the unit simplex. Then, we project this normalized 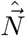 into the feasible region of the route *A*. The euclidean distance between this normalized 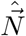 and its projection onto route *A*’s feasible region 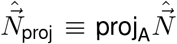 provides a heuristic for how likely 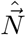 is to have been generated by route *A*. The farther away from the feasible region, the less likely. We use this distance to rank routes and discard routes by default with a feasible region greater than 0.2 away from the observed 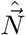.

#### Feasible route regions overlap

In the example above of the two routes for (3, 2) of *A*_major_first_ and *A*_minor_first_, each has its own constraints, which, after normalizing 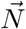 such that 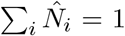, delineates a feasible region of valid multiplicity distributions for that route. If these regions were non-overlapping, then selecting the most likely route would be straightforward. However, the regions overlap – in fact 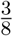 of *A*_major_first_’s region are shared and 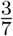 of *A*_minor_first_’s region are shared. Therefore, the two routes possess a non-identifiable region of intersection where both routes are consistent with the observed multiplicities. A second heuristic for selecting a route we employ is the inside margin – how far away from the boundary of each route is observed 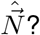 Under tie breaks of both routes being valid, we use this inside margin.

#### Clustering

We employed two types of clustering to detect cross-segment patterns in individual samples. The first type of clustering (time-point clustering) was performed in order to detect punctuated gain events that span multiple chromosomes, namely WGD and ccPG events. The second type (pooled-segment clustering) was performed in order to link segments of the same copy number, which may result from the same evolutionary route.

#### Time-point clustering – detecting punctuated gain events (WGD and ccPG)

In order to detect punctuated gain events, we compare, per-chromosome, the observed distribution of timing to a constant rate, under a Poisson distribution, in sliding windows of 0.06 units of molecular time. Per-chromosome p-values from the Poisson scan are corrected with Benjamini-Hochberg at *α* = 0.05. After scanning for peaks across each chromosome, chromosomal peaks are merged to detect sample-level timing events. Each merged event is then classified by its span across the genome. Events spanning either more than 40% of the genome that is sufficiently powered for timing or at least eight chromosomes were classified as WGD. An event meeting neither criterion for WGD but spanning at least two chromosomes is classified as a ccPG. Events on a single chromosome are labeled as chromosome-specific and are not counted as ccPGs.

#### Pooled-segment clustering – detecting segments with the same evolutionary history

For each copy-number state in a tumor sample, contiguous segments of that state are first merged into blocks. Blocks and remaining segments are then clustered by their multiplicity distributions with DBSCAN [46] (epsilon parameter 0.07, minimum cluster size 3) to find distinct pools of similar segments. The VAFs of SNVs within all segments of a cluster are then pooled to re-estimate the composite multiplicity distribution, and then re-time gains. These timing solutions then provide higher mutation count timing results. Segments within a pooled-segment cluster that have a low mutation count may be individually noisy, yet they still contribute informatively to a pooled-segment cluster. We time all amplified segments containing at least 10 SBS1/SBS5 mutations, with 80% of all the segment length in PCAWG data satisfying this condition (**Supplementary Figure S3a**). Pooling improves the accuracy of our timing estimates by increasing the count of mutations used in each route.

#### Benchmarking through simulated whole-genome tumors

We simulated whole-genome tumors using a custom script which simulates both WGD events and ccPG events, with a distribution of the number of SNVs per segment and SNV depth distribution based on the depth distribution of PCAWG tumors with a median depth of 50. Purity of tumor samples varied from 0.2-0.9. Using this model, we generated 3788 whole genome tumor simulations, with number of WGD events ranging from 0-2 and number of ccPG events ranging from 0-3. Focal and arm-level gains and losses follow Poisson distributions, with an elevated rate of loss following WGD.

We start by simulating the gain trajectory of each segment implied by the macro-events, e.g. WGD or ccPG – i.e. every segment will start as diploid, and then undergo changes in copy-number according to the tumor’s number of WGD or ccPG events and other more local gains and losses. This provides a clear evolutionary history for each segment, describing when each gain or loss occurred and on which allele. Therefore, per segment, we select a matrix that is consistent with the route implied by the simulation, and the time intervals between each event.

For instance, for a (3, 2) segment in a WGD simulated tumor, we might have 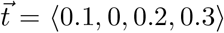, which, seeing as *t*_2_ = 0, would imply the first two gains are simultaneous, as we might find with a WGD event. Then, seeing as there are two possible routes to (3, 2), the simulation will pick one consistent with the path observed. Once the route *A* is picked, then multiplicities are simulated under a multinomial distribution generated from 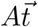. Then, each SNV’s VAF is simulated from a binomial distribution with parameters, like depth, based on the PCAWG cohort. The resulting VAF distribution is then used by Tau to estimate timing and enable comparison of estimation versus ground truth.

#### Segment-level benchmarking

Tau then produces per-segment estimates for the simulation, which can be directly compared to the true time. To quantify error in Tau’s estimation, we calculate the time of each gain per segment, i.e. the cumulative version of 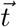, denoted 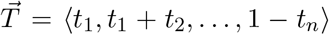. We compare the estimated gain times, 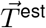 to the true times, 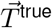 with the metric of maximum gain error: 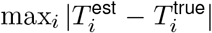, which provides a maximum on how poorly a segment has been timed (**Supplementary Figure S3b**). In the case of underdetermined gains, 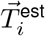 will have a range of values, rather than a single value, and so the closest value to 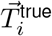 is used.

#### Event-level benchmarking

In addition to per-segment timing estimation, Tau clusters cross-segment timing results to infer the timing of larger (e.g. ccPG and WGD) events. Therefore, the output also produces per-event timing estimates to compare with the ground truth simulation (**Supplementary Figure S6a**). By running Tau on our simulated tumor samples, we benchmarked Tau’s ability to detect such events in different circumstances by matching Tau-estimated event times and ground truth event times with a tolerance of ±0.15 molecular time units. Unmatched inferred events were counted as false positives, and unmatched simulated events as false negatives. From this matching scheme, we calculated the median absolute error, along with recall and precision.

#### Time to whole-genome doubling

To characterize the timing of whole-genome doubling (WGD) across the cohort, we modeled the molecular time (on the standard 0–1 evolutionary timeline) to WGD acquisition as a time-to-event process with right-censoring. For each sample, the time of a WGD event was taken as its inferred molecular time; samples with no detected WGD were treated as right-censored at molecular time 1 (i.e. no WGD observed within the fully reconstructable timeline). We fit both single-parameter exponential (constant-hazard) and two-parameter Weibull time-to-event models by maximum likelihood; because Kaplan–Meier estimates showed a rising, non-constant hazard (low risk early in molecular time, increasing later) rather than the memoryless behavior implied by a constant-hazard process, and because the Weibull model achieved a substantially lower AIC, we report Weibull-based estimates throughout. To separate distinct questions about WGD dynamics, we fit four related but non-equivalent models: (i) time to first WGD across the entire cohort, with tumors that never acquire a WGD right-censored at molecular time 1, yielding a population-wide rate; (ii) time to first WGD restricted to samples that acquire at least one WGD (fully observed, with no censoring), which avoids assuming that WGD-negative tumors are simply “still at risk” rather than a biologically distinct population; (iii) a pooled-transition model treating every WGD-to-WGD interval within a sample (from tumor initiation to the first WGD, and from each subsequent WGD to the next, with the interval following a sample’s final observed WGD right-censored at 1) as an independent draw from a single shared process, to estimate an overall rate independent of whether a given WGD was a tumor’s first or a subsequent event; and (iv) a recurrence model isolating the interval between a first and second WGD specifically, conditioned on a first WGD having already occurred, to test whether recurrence hazard differs from the hazard of first occurrence. Model medians and 95% confidence intervals were obtained via the exact Poisson/chi-squared relationship for exponential fits, and via percentile bootstrap (resampling at the sample level and refitting each replicate) for Weibull fits, for which no closed-form censored-data interval exists. Comparisons between nested or overlapping estimates (e.g. WGD-positive-only versus entire-cohort medians, which share the same underlying samples) used paired bootstrap resampling rather than independent two-sample tests, since the compared quantities are not statistically independent. All models were fit both pooled across the cohort and separately within each tumor type with at least three WGD events.

### Gain heatmaps

#### Partitioning gains into WGD-linked, ccPG-linked, and unlinked categories

In order to classify gains as being associated with WGD events (WGD-linked) or ccPG events (ccPG-linked) or neither (unlinked), we looked at the distributions of gains around detected WGD events and ccPG events. Based on these distributions, we chose a threshold of ±0.2 for WGD events, and ±0.1 for ccPG events.

#### Normalization procedures for heatmaps

For the gain heatmaps, each gain event contributes total mass 1, spread as 1*/n*_solutions_ where *n*_solutions_ is the number of solutions – one for determined gains, but many more for underdetermined gains. This ensures that underdetermined gains do not contribute more but simply have their weight equally distributed across all consistent solutions. We then normalized each position in the heatmap by the total number of segments amplified at that genomic position, therefore calculating the number of gains per segment.

#### Oncogene annotation

Oncogenes and their positions were obtained from OncoKB [47], using hg19 coordinates.

### Analysis of burst amplifications

#### Defining burst amplifications

We defined a burst as greater than or equal to 3 consecutive gains occurring within a span of 0.05 molecular time units. For determined segments, this classification is straightforward because there is only one solution. For underdetermined segments, we distinguished routes in which every admissible solution contains a burst from those in which at least one admissible solution contains a burst (**Supplementary Figure S12b**). Therefore, in total, we had three categories of burst - determined, burst under every solution, and compatible with a burst. In all three categories, we observed enrichment at WGD (**Supplementary Figure S12c**).

#### Defining excess bursts versus WGD-explained bursts

In order to distinguish between burst amplifications that are fully explained by a WGD, versus those with a number of gains that exceeds the number explicable by a WGD, we calculated thresholds per copy-number state for the number of gains. For the major and minor copy-number (*M, m*), a single doubling event can, at most, cause ⌊*M/*2⌋ + ⌊*m/*2⌋ gains. We therefore count the number of gains in the burst *B*, and compare this number to *D* = ⌊*M/*2⌋ + ⌊*m/*2⌋. If *B > D*, we define this as an excess burst, as it exceeds the total possible number of gains from a single doubling event. If *B* ≤ *D*, we call this a WGD-explained burst. This definition is not dependent on distance from a WGD event time, but simply uses the conservative threshold on the number of gains to rule out bursts that could potentially be caused by a WGD. Therefore, a WGD-explained burst can potentially be far away from WGD, but because the number of gains is compatible with a WGD, we define it as WGD-explained.

#### Matched null simulations

We simulated two forms of null distributions.

First, in order to discern whether excess bursts would be detected by Tau as false positives, we matched the mutation counts and copy-number states of all high-copy-number PCAWG segments containing at least three gains, then used a flat Dirichlet distribution to sample gain times and simulated SNV VAFs for each segment, from which multiplicity counts were estimated and timed exactly as for real segments. We completed this procedure ten times per segment. We then calculated the proportion of segments in the simulated null that met our criteria for definition of an excess burst (**Supplementary Figure S12a**).

Secondly, in order to test the association of excess bursts with WGD, we permuted WGD times randomly 20,000 times between samples and observed the proportion of excess bursts that were ±0.1 molecular time units away from a WGD. We then compared these proportions to the proportions of excess bursts ±0.1 molecular time units away from a WGD in the actual data (**Figure 6b, h**; **Supplementary Figure S12c**).

#### Excess burst hotspots

For each chromosome arm, we counted the samples carrying at least one excess burst within the arm and compared this to the number of samples without an excess burst on that arm. Then, we counted the samples carrying at least one excess burst versus no excess bursts on all other arms, and used these values to calculate an odds ratio per chromosome arm and a 95% confidence interval from the standard error of the log odds ratio. Arms with fewer than 5 excess burst carrier samples were excluded.

#### Estimating post-WGD loss

To estimate post-WGD loss across the genome, we looked at PCAWG samples with at least one detected WGD. We inspected the best-fitting routes for each segment and the estimated WGD time. Based on the number of copies before WGD, we inferred the expected number of copies per allele after WGD. For instance, if there is 1 copy of the major allele and 1 on the minor, then the expected number after doubling would be 2 on the major allele and 2 on the minor. We then compared this expected number to the actual number, to infer loss. For instance, we might observe 2 on the major allele and 1 on the minor allele after doubling, implying a total loss of 1 (**Supplementary Figure S9a**,**c**). As a conservative estimate, we used ±0.2 to link gains to a WGD event.

#### PCAWG cohort

We ran Tau on 2,703 tumor samples with PCAWG consensus SNV, allele-specific copy-number and purity calls, and MuSiCal signature exposures. Gain and segment-level analyses (**Figure 5, Figure 6, Supplementary Figure S8**–**Supplementary Figure S12**) use all samples. Sample-level prevalence and timing statistics (**Figure 3d–f**; **Figure 4, Supplementary Figure S7**) use one sample per donor and exclude seven prostate donors represented only by multiple metastases.

## Data and Code Availability

The Tau source code and analysis pipelines are available at https://github.com/gulhanlab/Tau/tree/main. The results of PCAWG analysis are presented as an interactive website at https://tau.sigscape.org/.

## Author Contributions

JB implemented the model, performed the analyses, prepared the figures, and co-wrote the manuscript. PJP provided conceptual guidance, contributed to the interpretation and framing of the results, and reviewed and edited the manuscript. DCG conceived and supervised the study, developed an initial prototype of the model, contributed to the analyses and figure design, and co-wrote the manuscript.

## Acknowledgements

This work was supported by grants to PJP (NIH R01CA269805, R01HG012573, and UM1DA058230) and DCG (DoD OC230088). We thank Christophe Boetto, Cait Harrigan, Jan Hummel, and Ignacio Vázquez-García for helpful comments and discussions that improved the manuscript, and Shannon Ehmsen for her input on the figure and logo design.

---

**Supplementary Figure S1:**
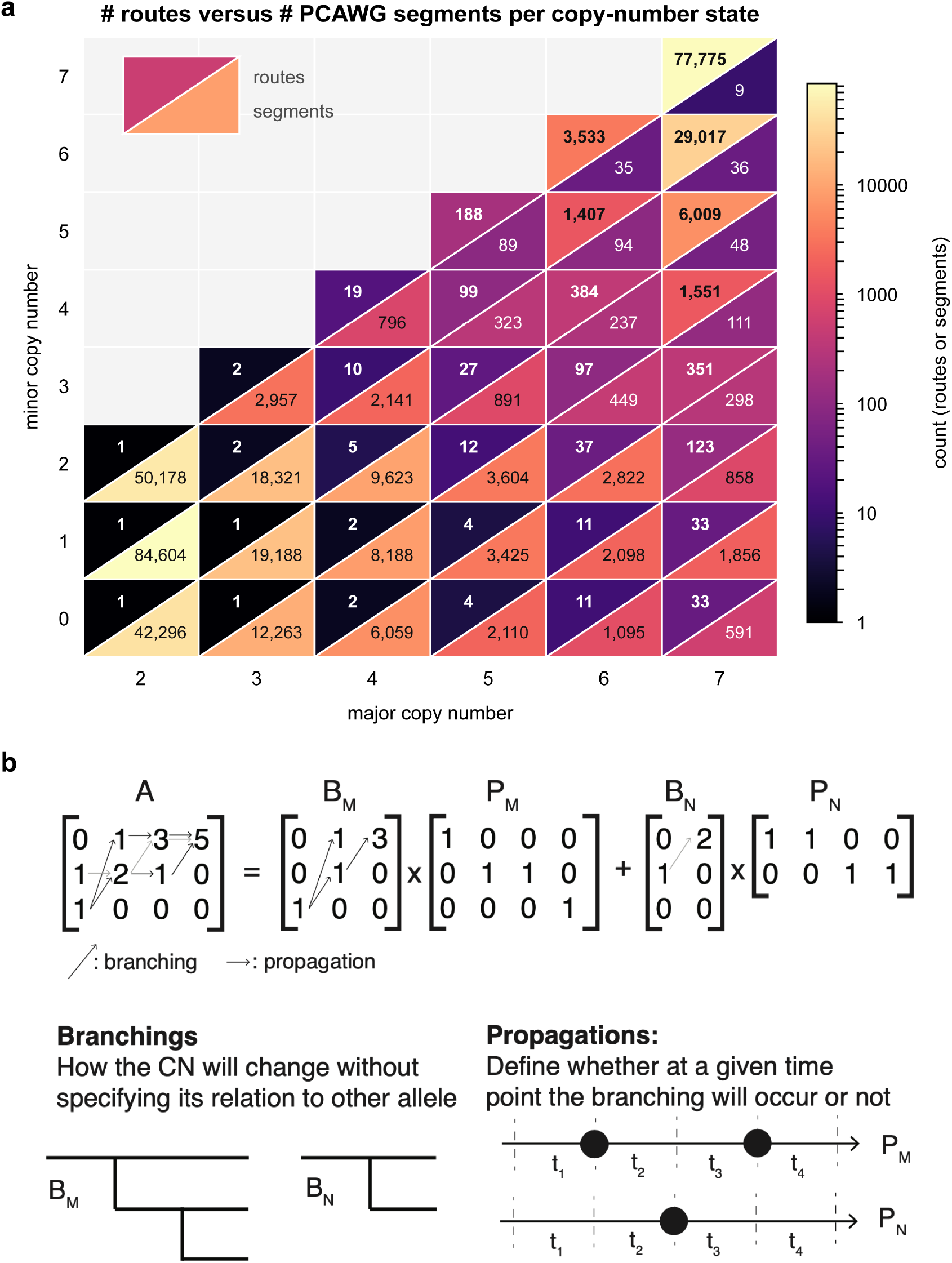
Growth of route complexity with copy number. **a**, Number of distinct gain routes and number of PCAWG segments observed, for each allele-specific copy-number state. Each cell is split: one half gives the route count, the other the segment count, on a shared logarithmic scale. Route number grows steeply with both major and minor copy number, while observed segments concentrate in low-copy states. **b**, Construction of route matrices: a route (*A*) to a state is decomposed into allele-specific (*M* = major, *N* = minor) routes (*B*_*M*_ × *P*_*M*_ and *B*_*N*_ × *P*_*N*_), and further broken down into branching (*B*_*M*_ and *B*_*N*_) and propagation (*P*_*M*_ and *P*_*N*_) matrices. Branching matrices specify how copy-number of SNVs on the M or N allele changes with new amplifications on the allele, while propagation matrices specify when these allele-specific amplifications will occur, relative to the other allele. This decomposition allows recursive construction of routes to any given copy-number state.

**Supplementary Figure S2:**
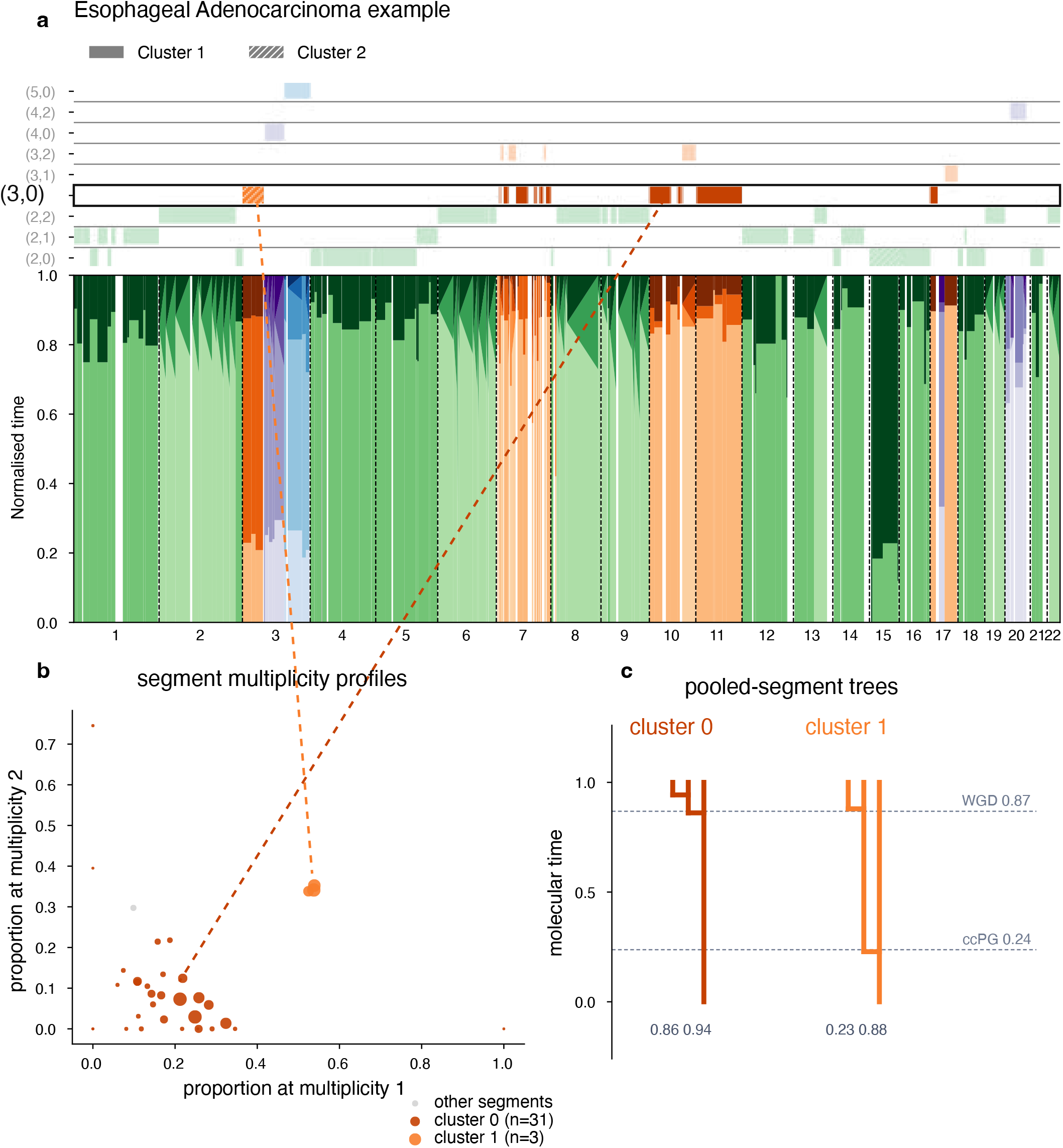
Pooled-segment clustering of multiplicity profiles. **a**, Esophageal adenocarcinoma example, with two (3, 0) pooled-segment clusters highlighted. **b**, Multiplicity profiles of individual segments of copy-number state (3, 0) in the esophageal adenocarcinoma, proportion of multiplicity 1 on x-axis and multiplicity 2 on y-axis. Two (3, 0) clusters shown, with lines drawn to show, in **a**, genomic position of clusters. **c**, Tree diagrams of the two (3, 0) pooled-segment clusters, with detected WGD (whole-genome duplication) and ccPG (cross-chromosomal punctuated gain) times annotated

**Supplementary Figure S3:**
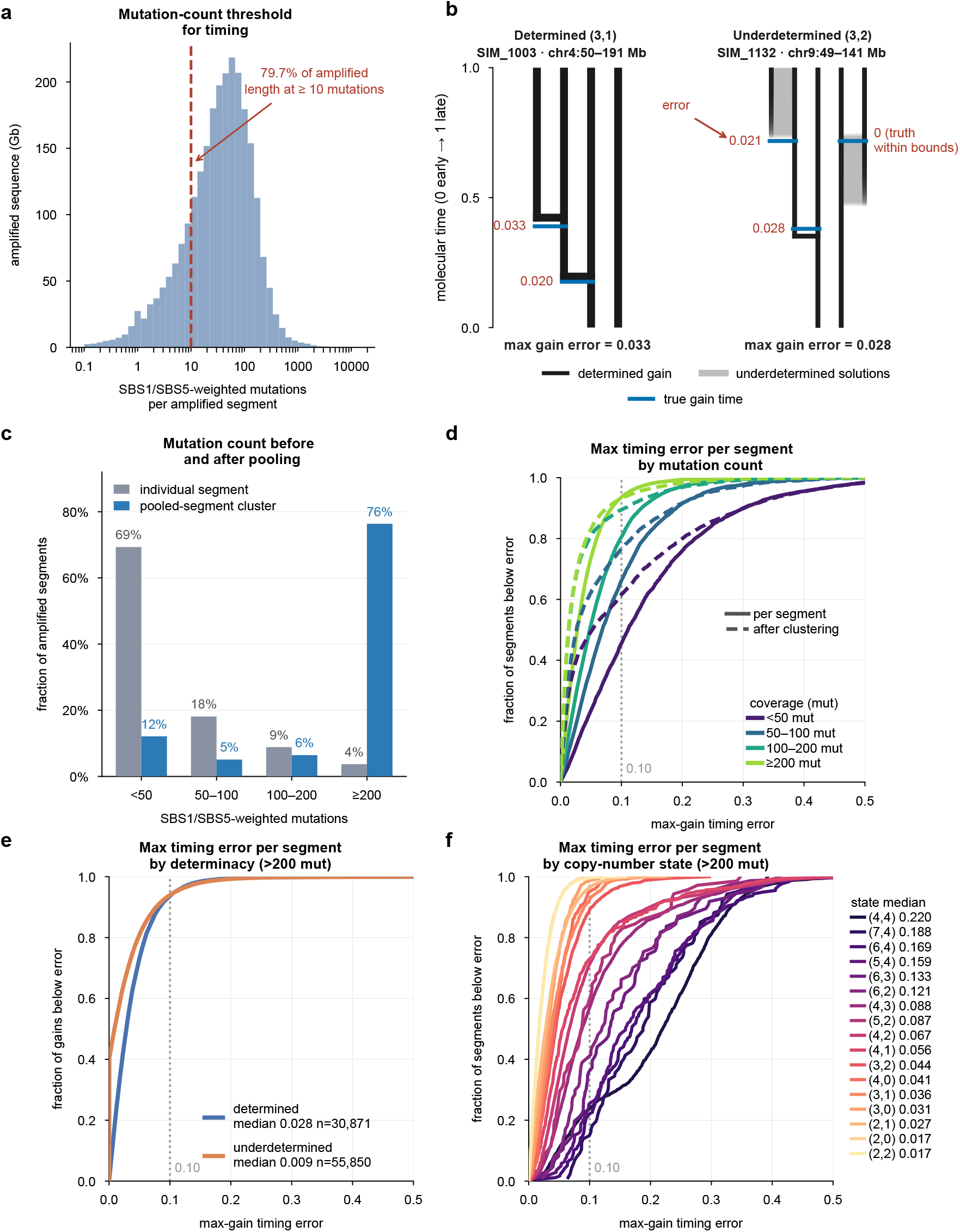
Segment-level benchmarking. **a**, Distribution of SBS1/SBS5-weighted mutation counts per amplified segment in PCAWG; dashed line, the threshold of 10 mutations below which segments are not timed. **b**, Definition of the error metric: the maximum absolute difference between estimated and true cumulative gain times across a segment’s gains. For underdetermined gains the closest point in the solution range is used. **c**, Fraction of PCAWG segments exceeding 200 weighted mutations, as individual segments and after pooling into route clusters. **d–f**, Cumulative distribution of maximum gain error in simulated tumors, stratified by mutation count before and after pooling (**d**), by whether the route is determined or underdetermined (**e**), and by copy-number state (**f**).

**Supplementary Figure S4:**
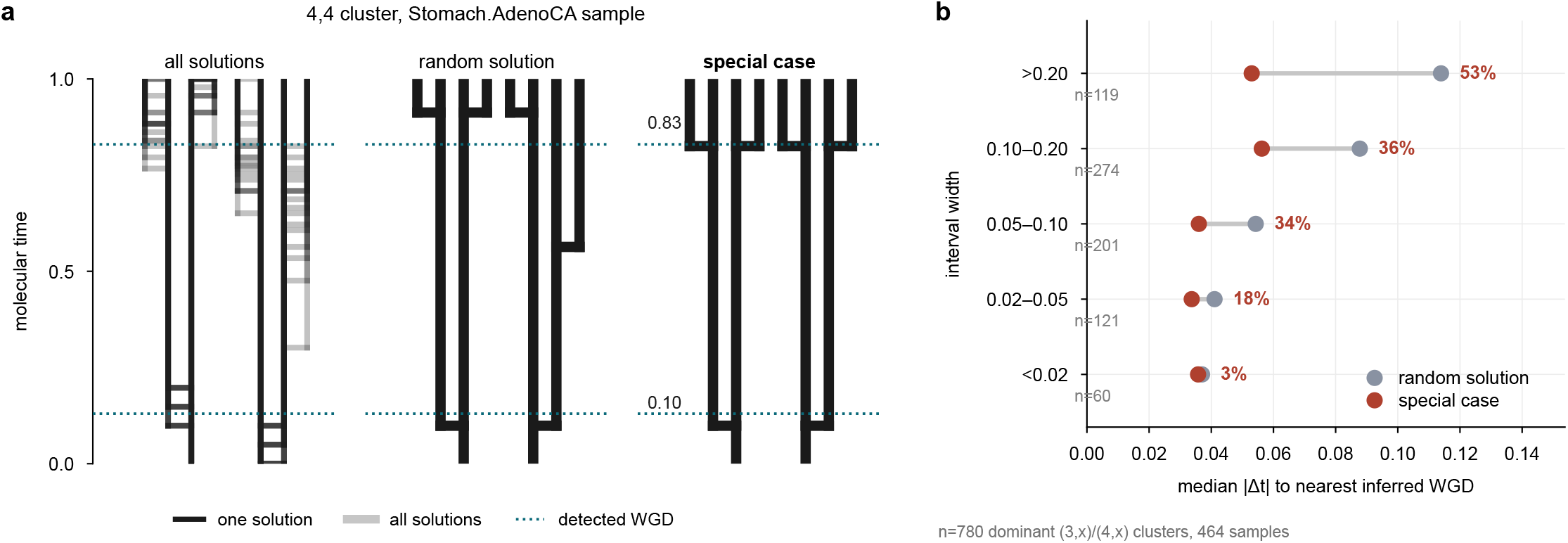
Underdetermined routes and special-case solutions. **a**, A single well-powered (4, 4) pooled-segment cluster from the double-WGD stomach tumor, drawn three ways: all solutions of its route overlaid, one arbitrary solution, and the special-case solution in which gains coincide on multiple branches. Detected WGD times are drawn for reference and do not enter the selection. **b**, Distance from the co-timed gains to the nearest inferred WGD, comparing an arbitrary solution with the special case, binned by the width of the solution range. The special case places gains closer to the WGD, and increasingly so where the range is wider.

**Supplementary Figure S5:**
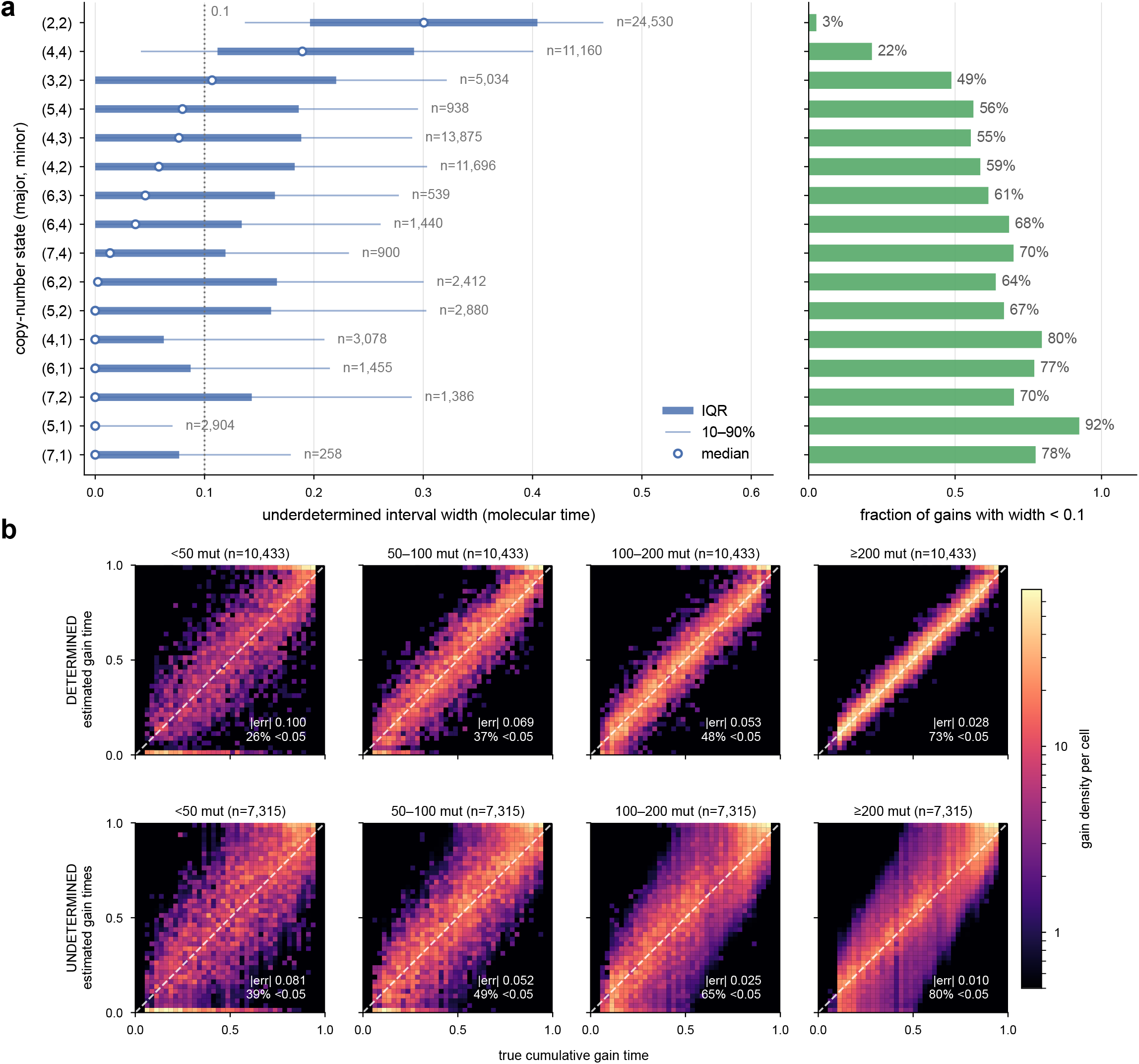
Range of possible solutions by copy-number state. **a**, Width of the solution range for each gain of an underdetermined route, by copy-number state (median, interquartile range and 10–90% interval). Several states are pinned to a point despite being underdetermined; (2, 2) and (4, 4) have the widest ranges. **b**, Estimated against true cumulative gain time in simulated tumors, binned by mutation count, shown separately for determined and underdetermined routes. The width of the solution range does not narrow as mutation count increases.

**Supplementary Figure S6:**
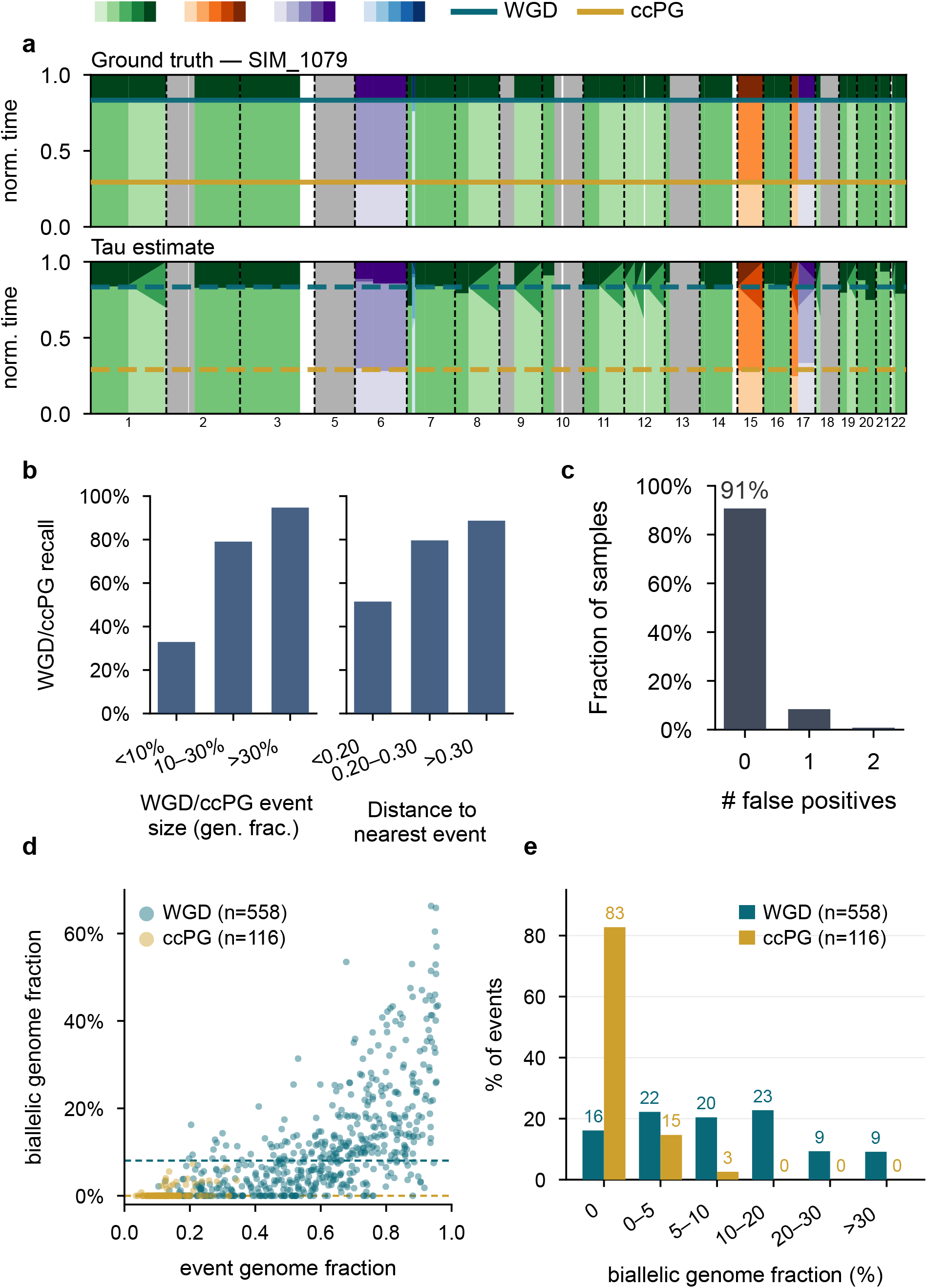
Benchmarking of WGD and ccPG detection. **a**, A simulated tumor with its ground-truth events alongside the events recovered by Tau. **b**, Detection recall as a function of event size and of the separation from the nearest other event. **c**, Number of false-positive events called per simulated sample. **d**, Biallelic genome fraction of detected event in PCAWG, against the fraction of the genome the event covers, for WGD and ccPG. **e**, Distributions of biallelic genome fractions of WGD and ccPG events.

**Supplementary Figure S7:**
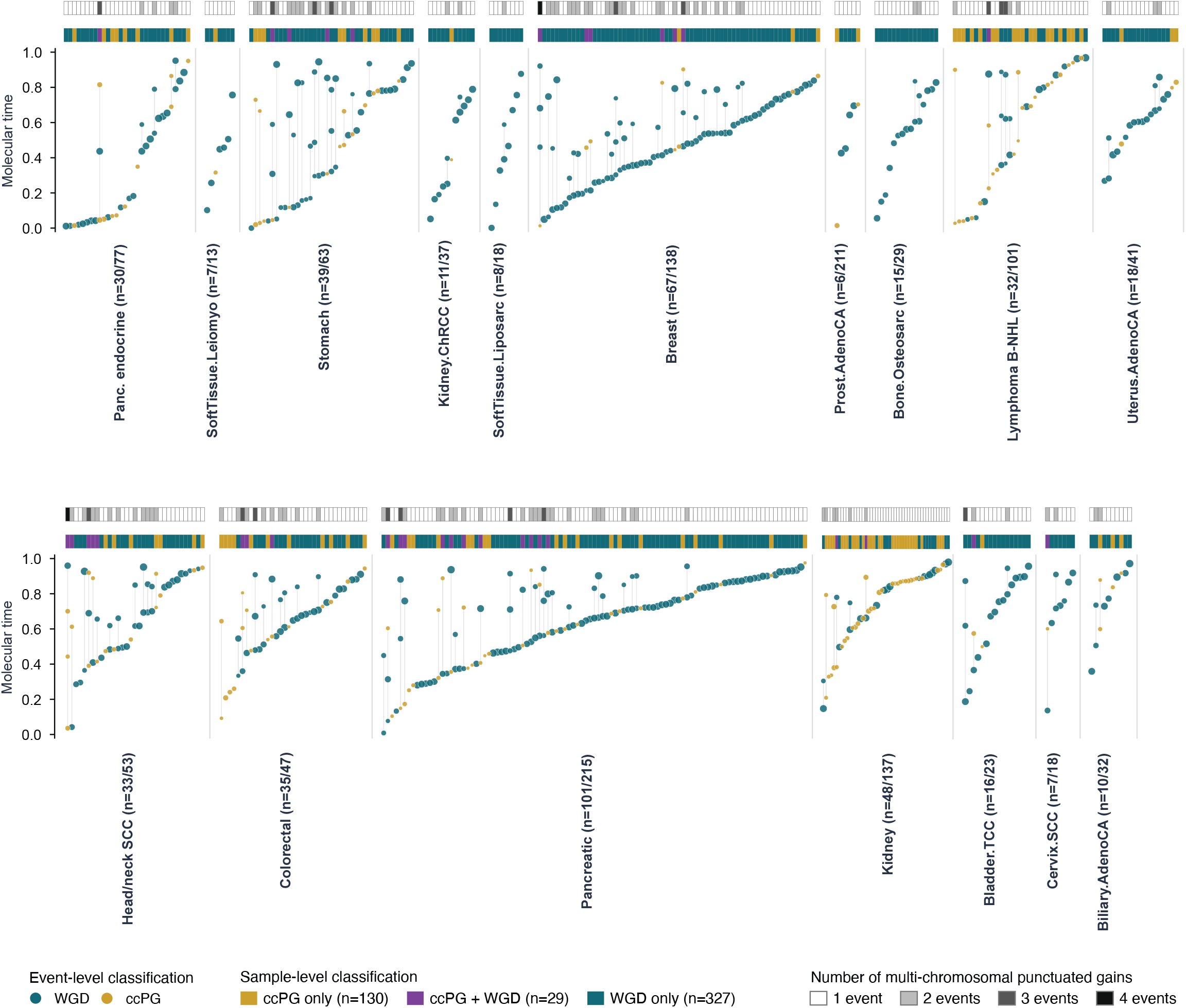
Per-sample WGD and ccPG event times in the remaining tumor types. As in Figure 4a.

**Supplementary Figure S8:**
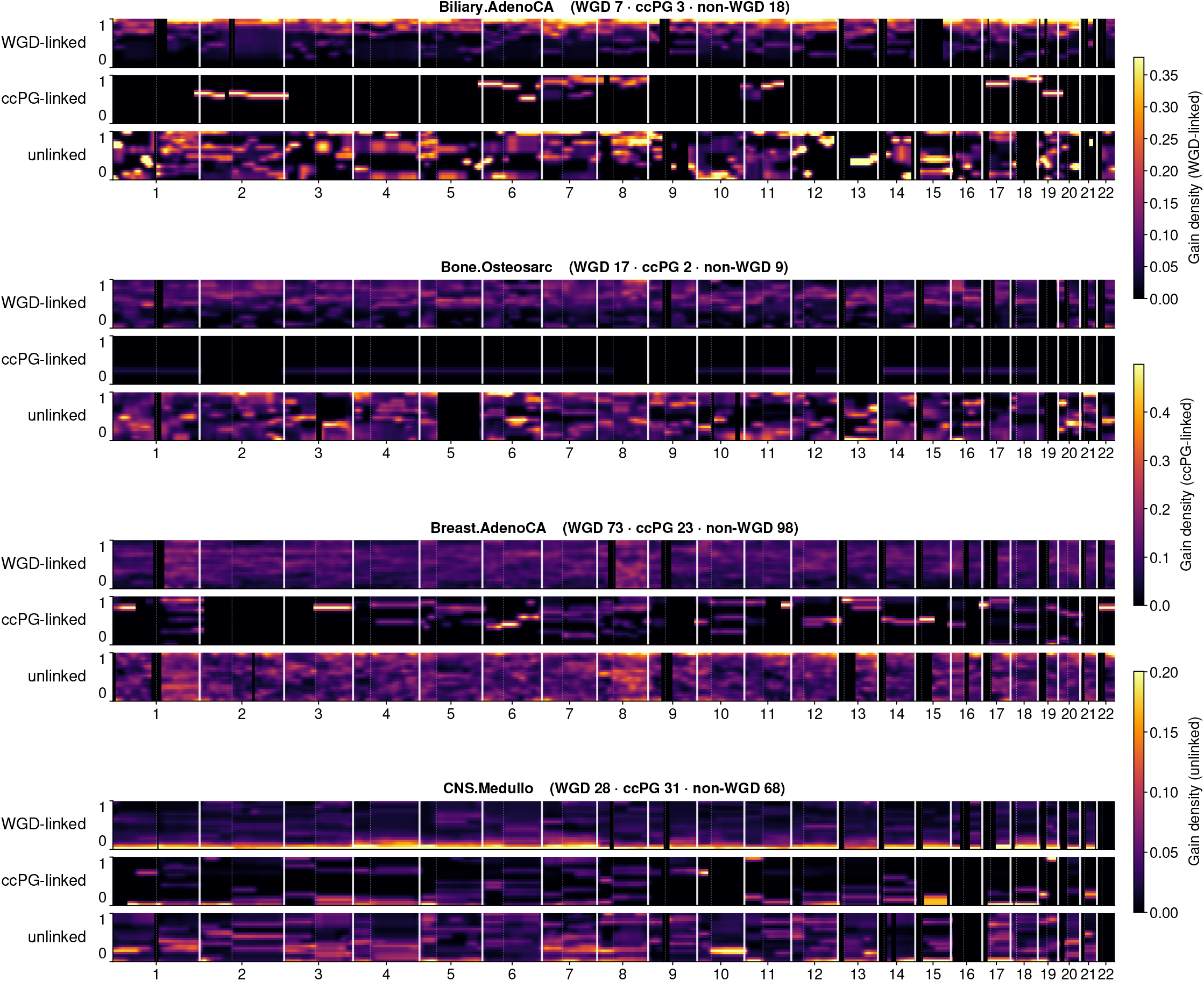
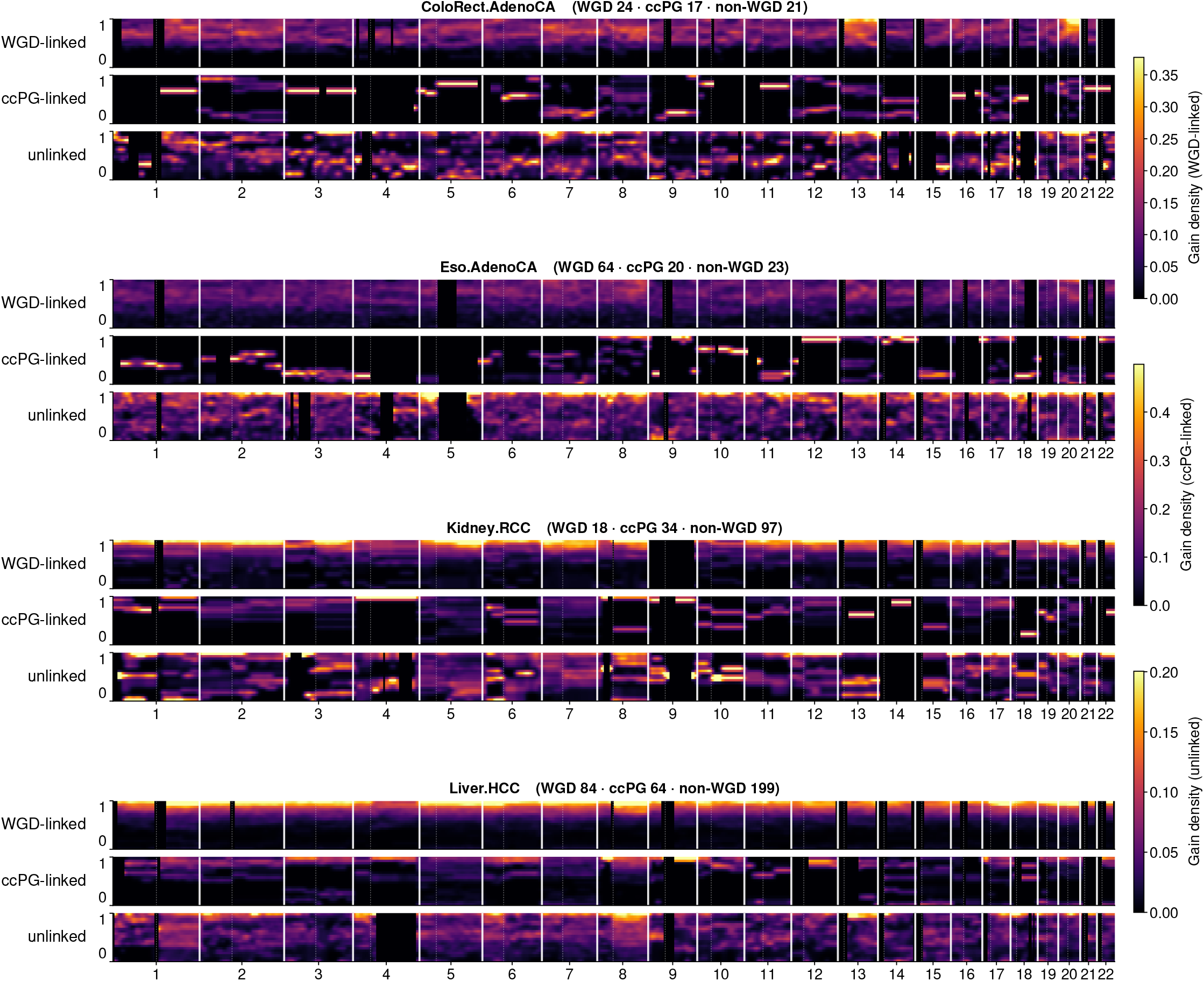
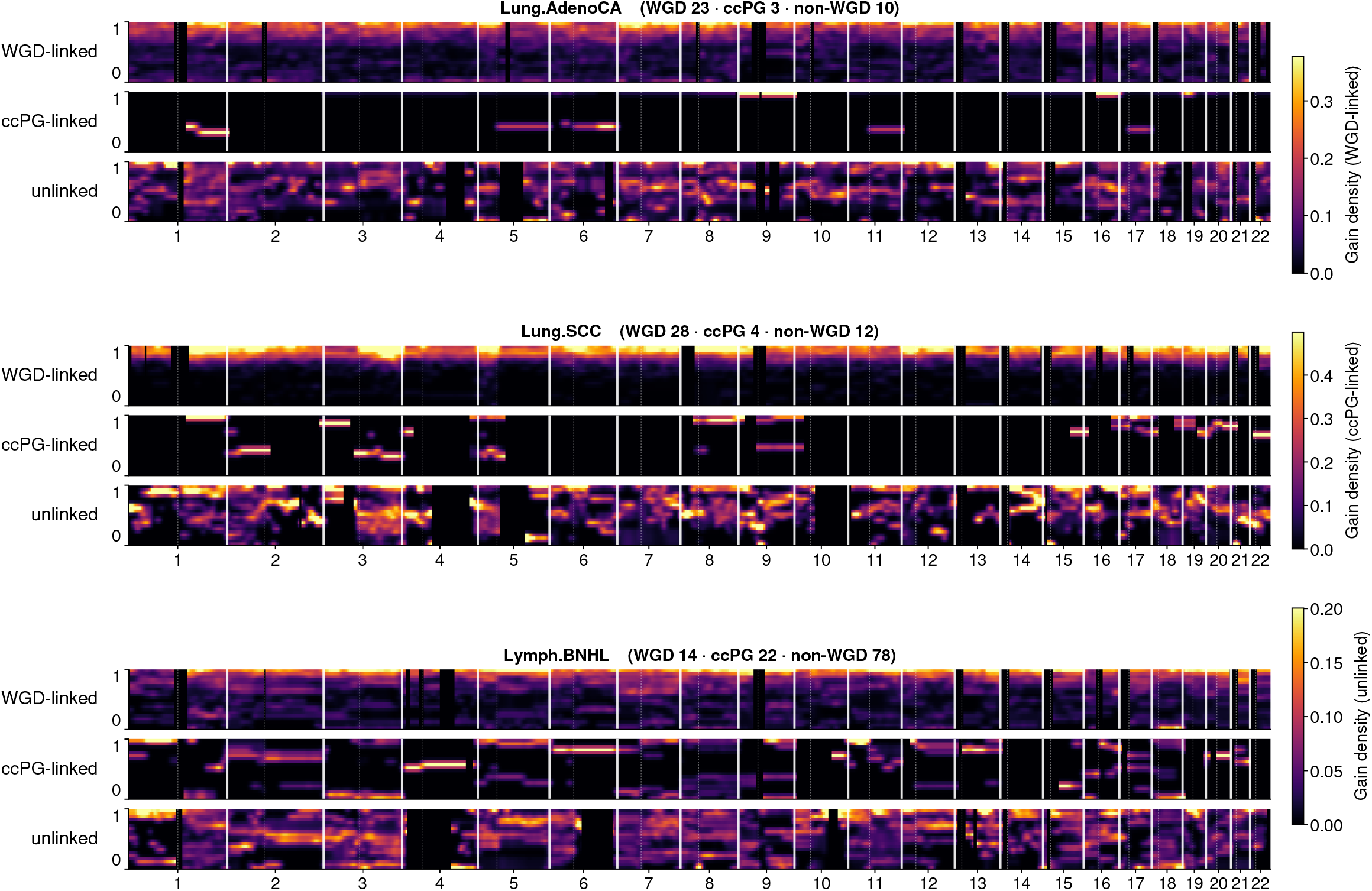
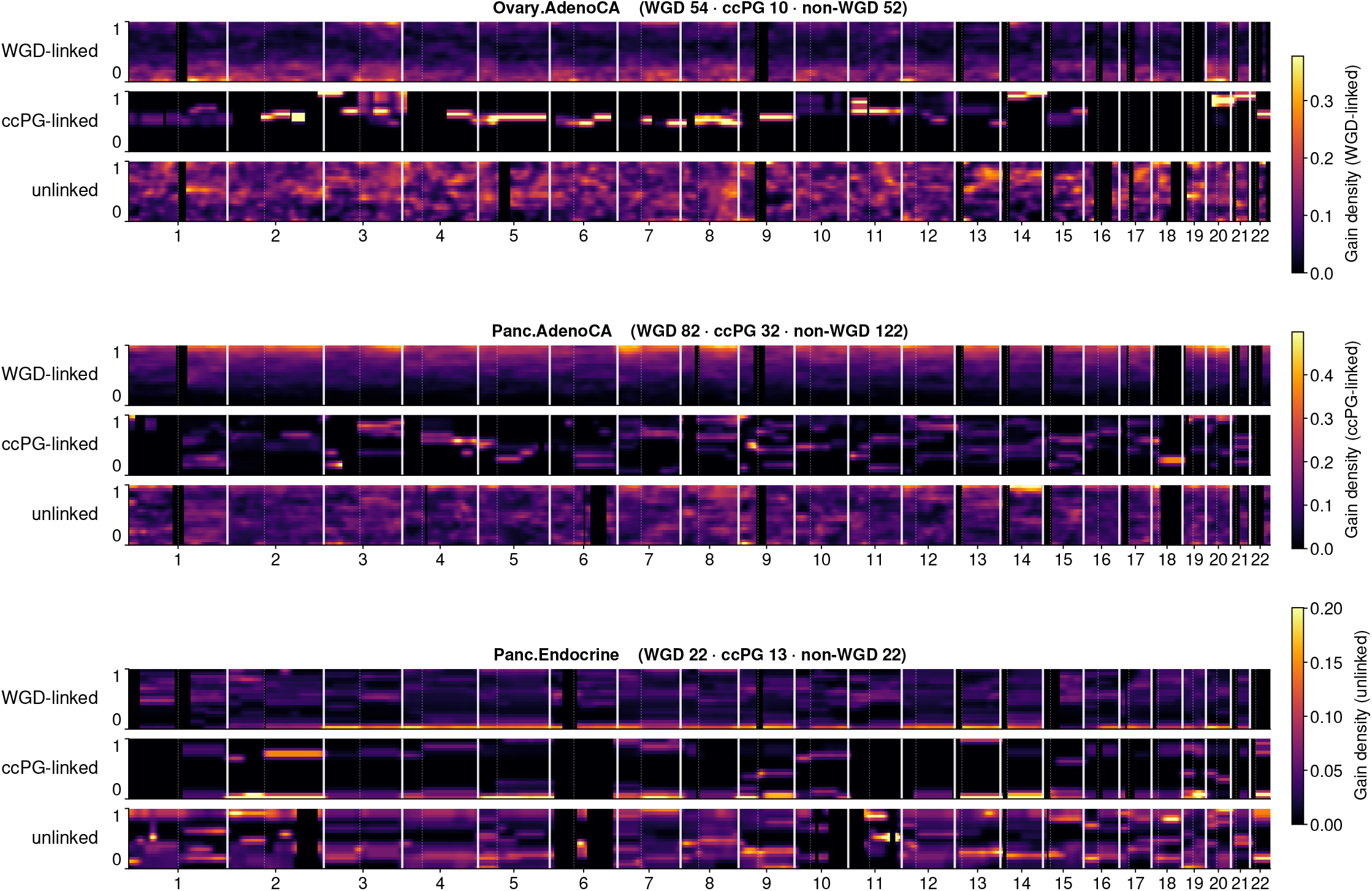
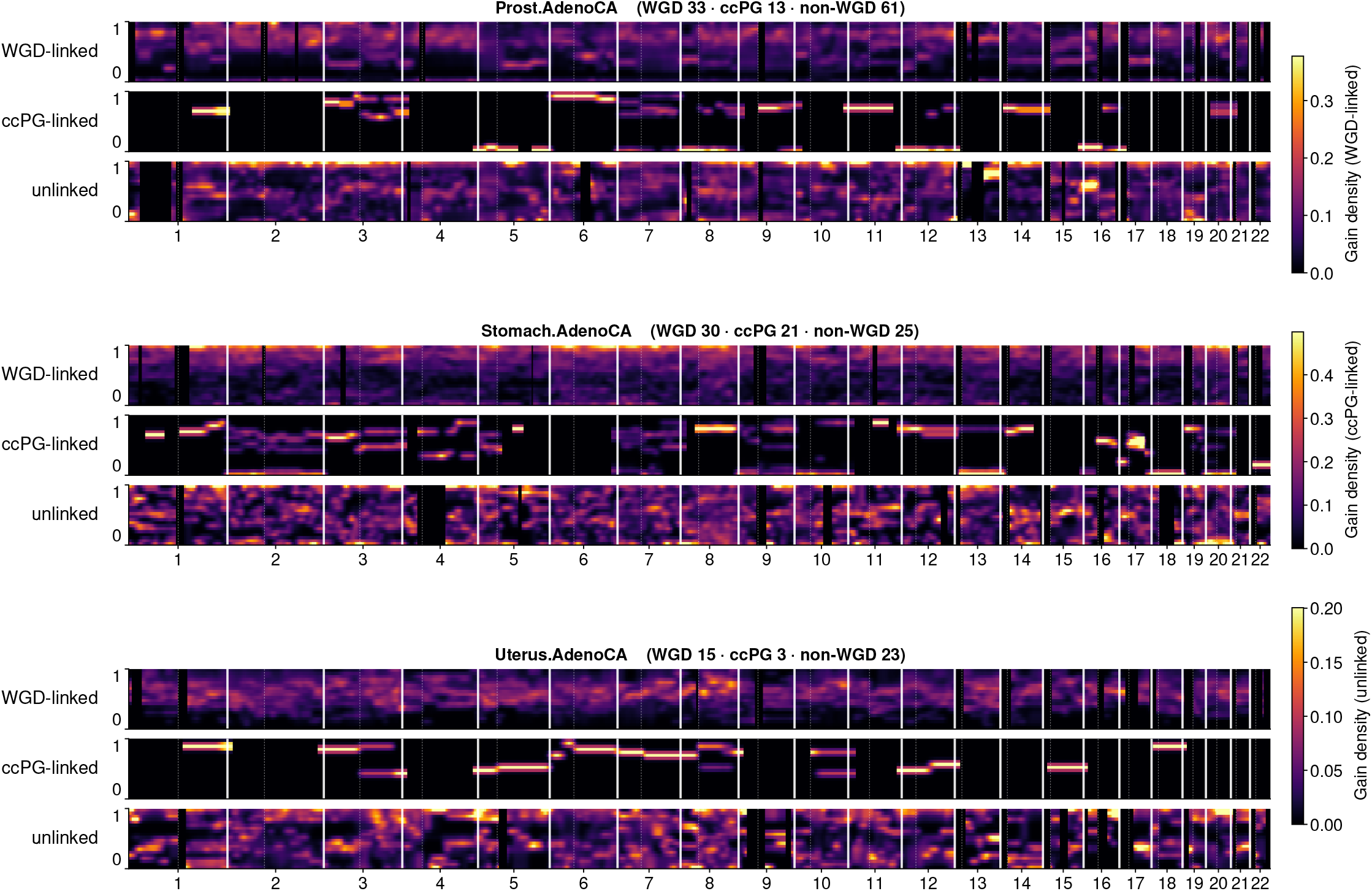
Landscape of gains in more tumor types. Genome position (*x*) against molecular time (*y*) for every tumor type with at least 25 timed samples and at least 5 WGD samples. For each type, gains are partitioned into three layers (rows): WGD-linked (within 0.20 molecular time of a detected WGD), ccPG-linked (within 0.10 of a detected ccPG and not already WGD-linked), and unlinked (neither). Each layer is divided by its own per-locus segment denominator, so color is gains per contributing segment. The color scale is shared across all five pages of this figure, so panels are directly comparable between pages, but is not shared between layers. Sample counts per type are given in each header. Biliary.AdenoCA, Bone.Osteosarc, Breast.AdenoCA, CNS.Medullo ColoRect.AdenoCA, Eso.AdenoCA, Kidney.RCC, Liver.HCC. Lung.AdenoCA, Lung.SCC, Lymph.BNHL. Ovary.AdenoCA, Panc.AdenoCA, Panc.Endocrine. Prost.AdenoCA, Stomach.AdenoCA, Uterus.AdenoCA.

**Supplementary Figure S9:**
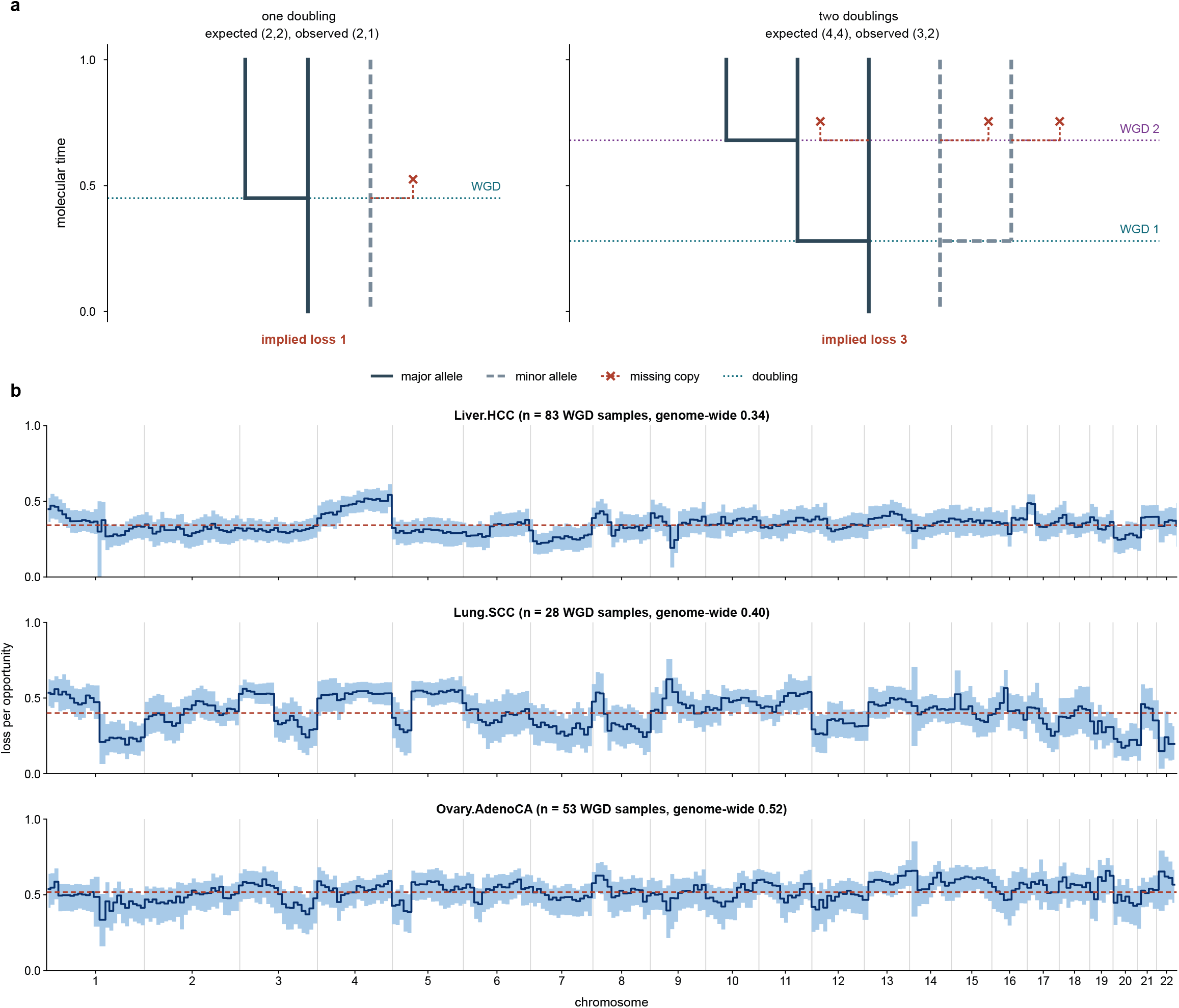
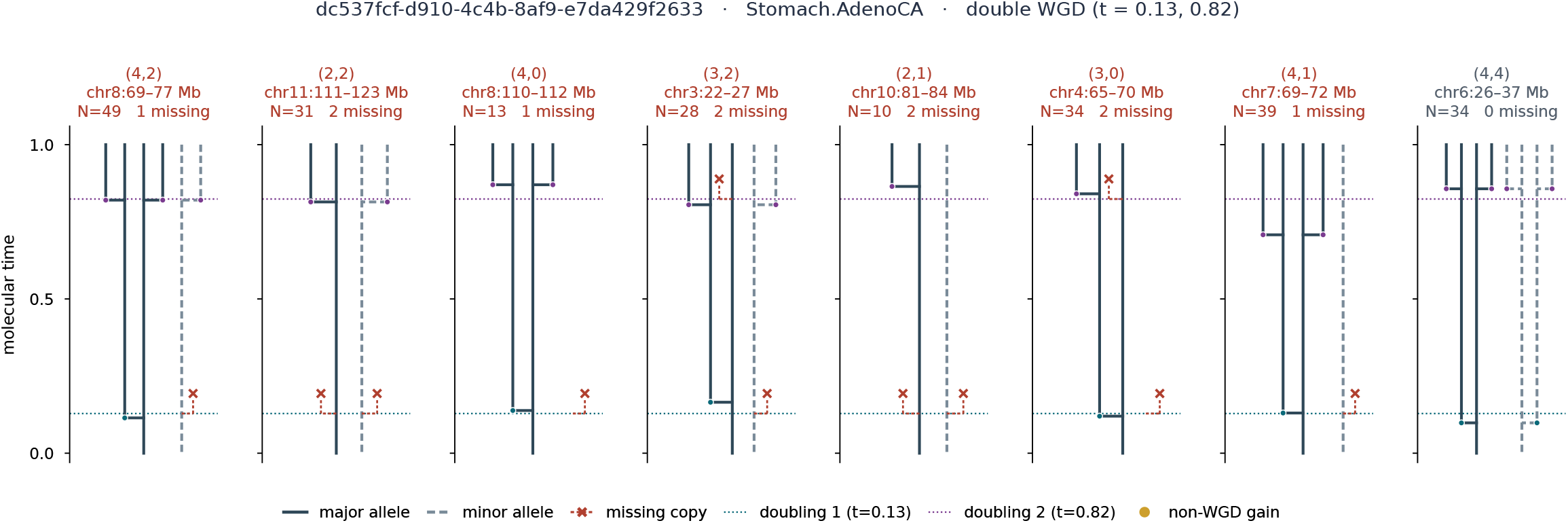
Quantification of loss across the genome. **a**, Schematic showing method for quantification of loss. Losses are inferred by inspecting the expected number of copies following WGD (left), or double WGD (right) compared to the observed distribution of WGD-linked gains and final copy number. **b**, Loss across the genome in hepatocellular carcinoma (Liver.HCC), squamous cell lung carcinoma (Lung.SCC), and ovarian adenocarcinoma (Ovary.AdenoCA), normalized by number of gains to lose at WGD (loss per opportunity). **c**, examples of loss calculations from a double-WGD stomach adenocarcinoma.

**Supplementary Figure S10:**
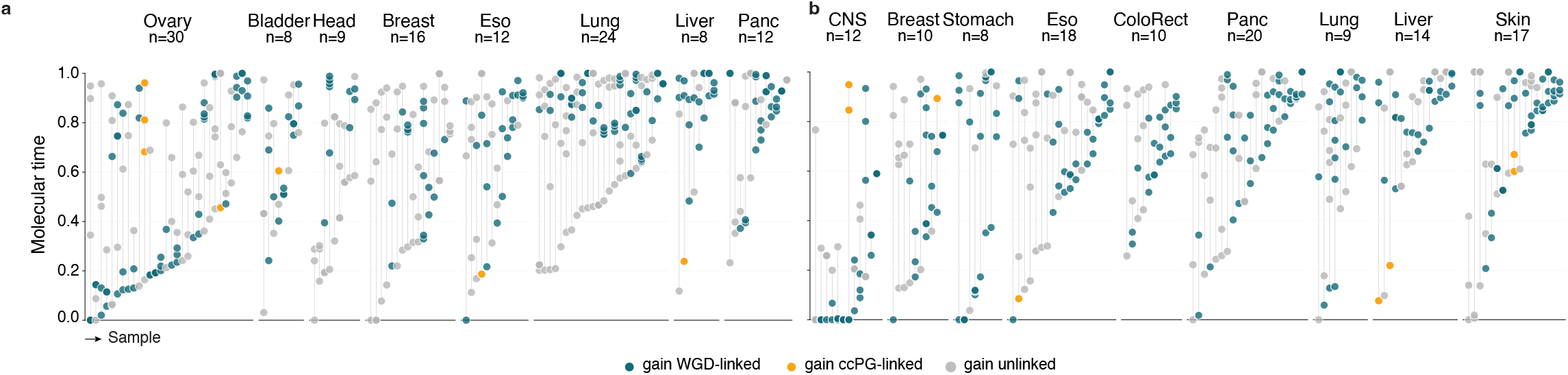
Oncogene amplification patterns: sequence of gains at *PIK3CA* (**a**) and *EGFR* (**b**), by tumor type. Each column is a sample; points give the molecular time of every gain at that locus, colored by whether the gain is WGD-linked (blue), ccPG-linked (gold) or unlinked (grey). Samples are ordered by the time of their first gain. Gains dispersed across long intervals indicate gradual amplification, whereas points stacked within a narrow interval indicate a burst.

**Supplementary Figure S11:**
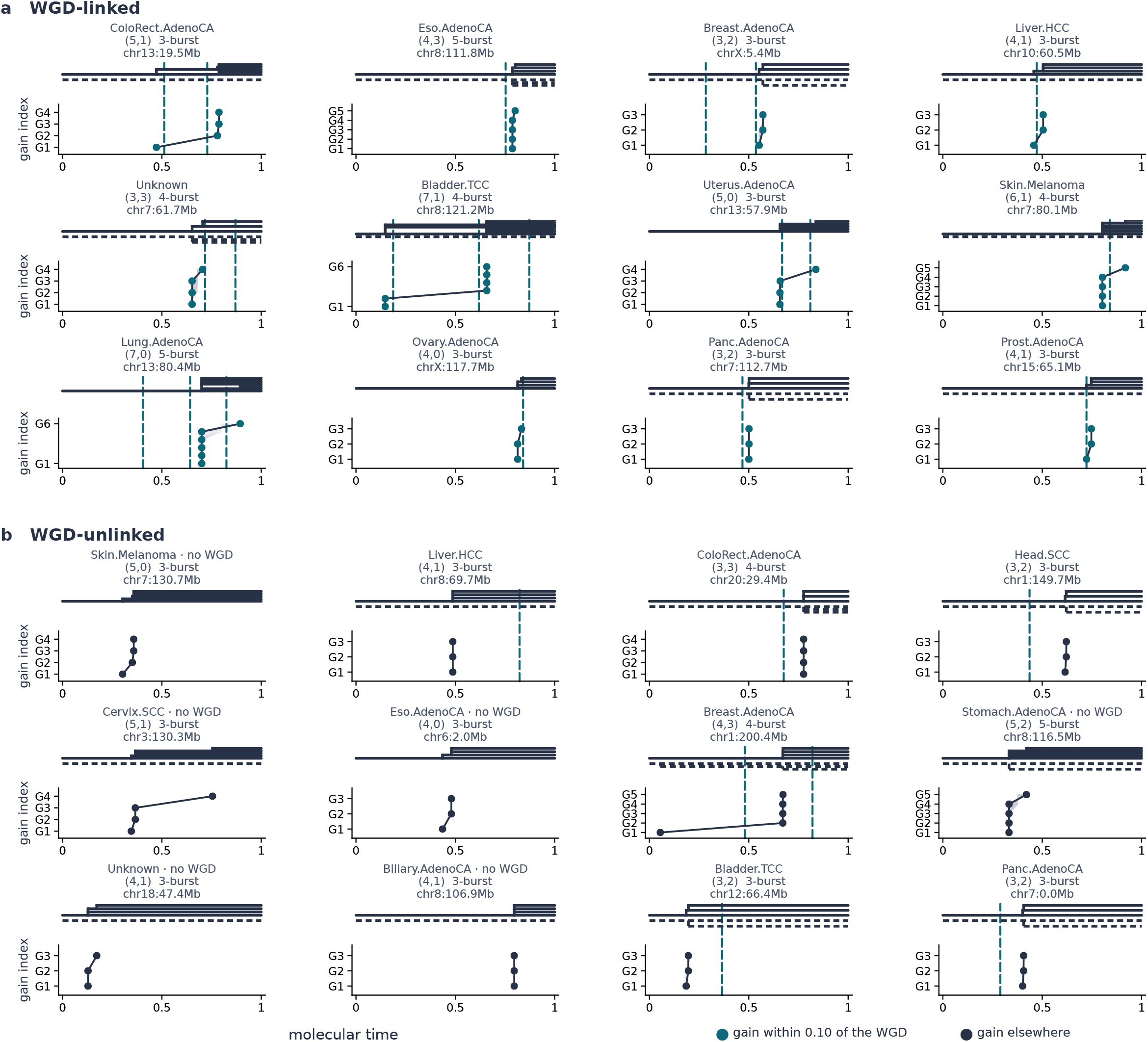
Examples of burst amplifications across the PCAWG cohort. Each panel shows an individual burst amplification from some sample or tumor type.

**Supplementary Figure S12:**
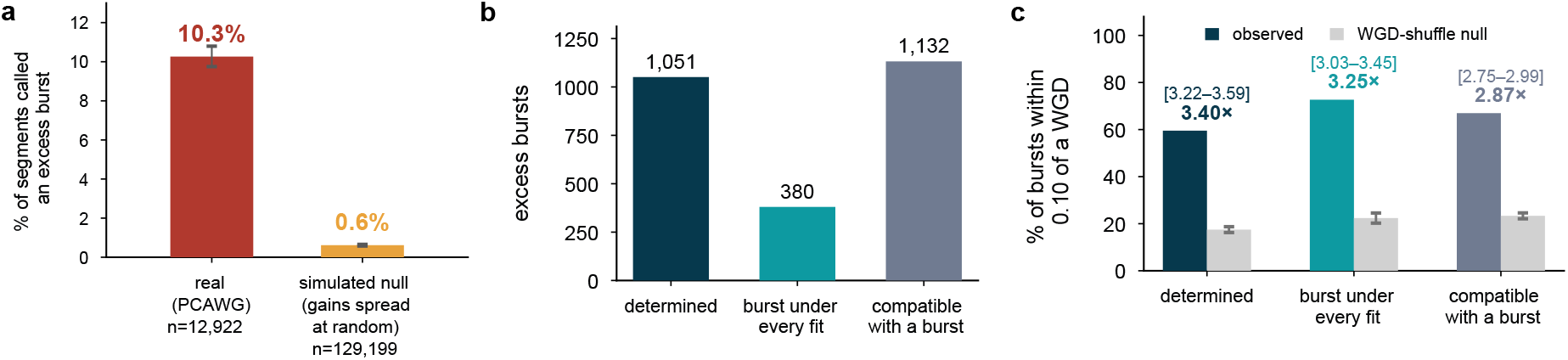
Excess burst amplifications exceed a matched null and show enrichment at WGD across both determined and underdetermined segments. **a**, Bar chart showing percentage of Tau-estimated excess bursts in null simulations with gains spread at random matched to the distribution of copy-number states versus percentage detected in actual PCAWG segments. **b–c**, Counts (**b**) and WGD enrichment (**c**) of excess bursts for determined segments (left), underdetermined segments with a burst detected in every solution (middle), and under-determined segments with a burst in only some solution, i.e. compatible with a burst (right)

**Supplementary Figure S13:**
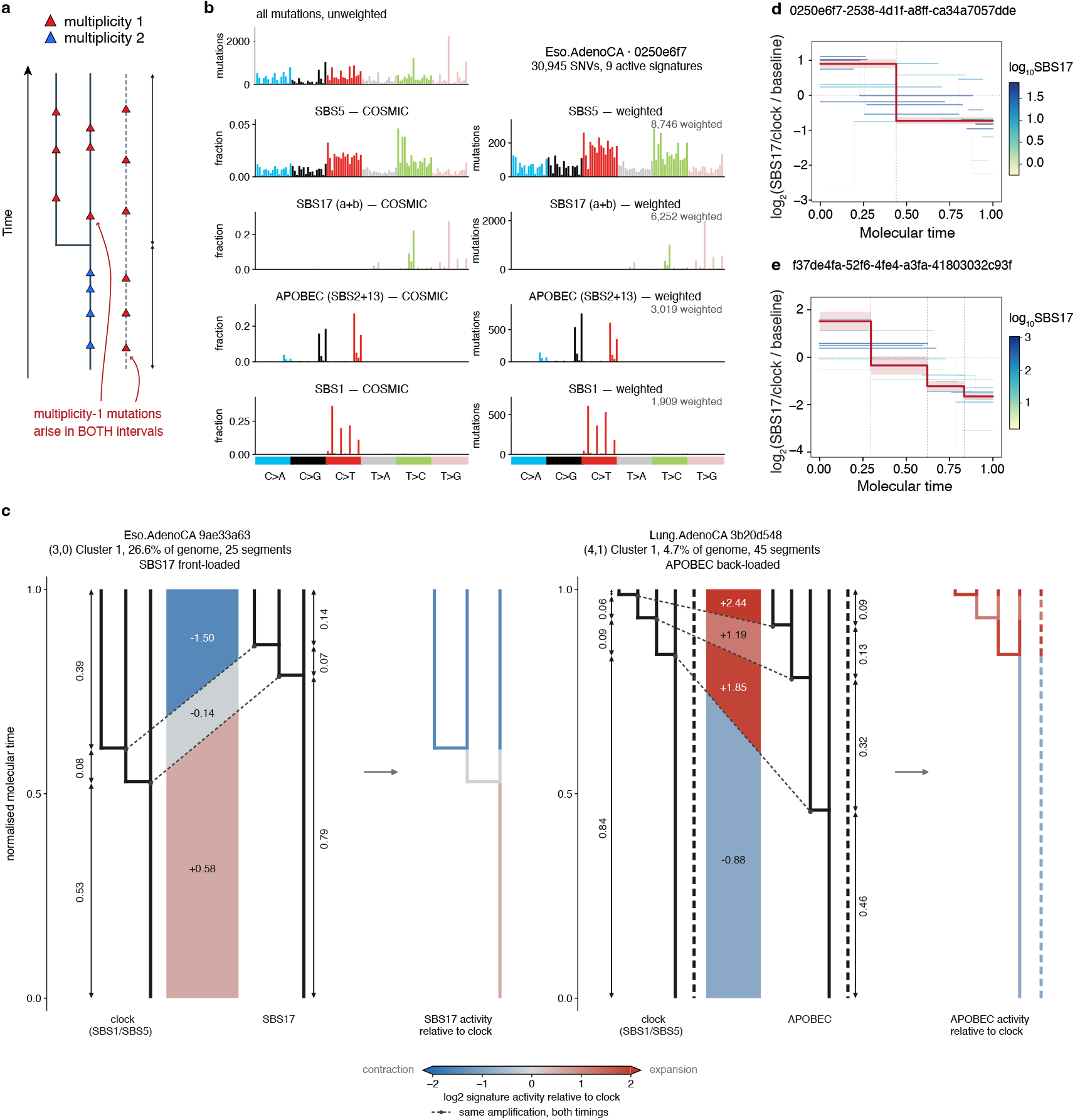
Timing different mutational process activity relative to clock-like mutations. **a**, Schematic showing how multiplicity 1 mutations can arise in different time intervals between gains. Mutations marked as triangles (red = multiplicity 1, blue = multiplicity 2), major allele solid line, and minor allele dashed. **b**, Weighting mutations by probability of coming from different mutational signatures in order to time different mutational processes. On first row is the unweighted spectrum of mutations from an esophageal adenocarcinoma PCAWG tumor. Below, left column shows COSMIC spectra for different mutational signatures; right shows distribution of tumor’s mutation weights for corresponding COSMIC signature, each showing high similarity to the COSMIC reference. **c**, Two examples of timing mutational processes relative to SBS1+SBS5 timing results. First tree in each example shows the SBS1+SBS5 timing. Colored intervals show relative size of intervals (red = expansion, i.e. higher mutational process activity relative to SBS1+SBS5, blue = contraction, i.e. lower mutational process activity). Tree to the right of the colored intervals shows timing with mutations weighted by mutational process. Third tree shows SBS1+SBS5 tree colored by expansion or contraction of mutational process in each interval. SBS17 (left) in esophageal adenocarcinoma shows higher activity earlier (front-loaded), with SBS17 tree showing larger intervals (expansion) early and smaller intervals (contraction) late, relative to SBS1+SBS5 tree. APOBEC (right) in lung adenocarcinoma shows opposite effect, with higher activity later (back-loaded), with APOBEC tree showing larger intervals (expansion) late and smaller intervals (contraction) early, relative to SBS1+SBS5 tree. **d**,**e**, The stepwise-transition model of signature activity, fitted to two samples for SBS17 activity, is shown as a red step function with its standard error (red band around line); in **d**, BIC selected a model with two activity states. In **e**, it selected a model with four activity states. Intervals contributing to the fit from all clustered routes are shown as lines, each spanning certain window in molecular time, and colored by SBS17 activity during that interval.

